# Long-read sequencing and genome assembly of natural history collection samples and challenging specimens

**DOI:** 10.1101/2024.03.04.583385

**Authors:** Bernhard Bein, Ioannis Chrysostomakis, Larissa S. Arantes, Tom Brown, Charlotte Gerheim, Tilman Schell, Clement Schneider, Evgeny Leushkin, Zeyuan Chen, Julia Sigwart, Vanessa Gonzalez, Nur Leena W.S. Wong, Fabricio R. Santos, Mozes P. K. Blom, Frieder Mayer, Camila J. Mazzoni, Astrid Böhne, Sylke Winkler, Carola Greve, Michael Hiller

## Abstract

Museum collections harbor millions of samples, largely unutilized for long-read sequencing. Here, we use ethanol-preserved samples containing kilobase-sized DNA to show that amplification-free protocols can yield contiguous genome assemblies. Additionally, using a modified amplification-based protocol, employing an alternative polymerase to overcome PCR bias, we assembled the 3.1 Gb maned sloth genome, surpassing the previous 500 Mb protocol size limit. Our protocol also improves assemblies of other difficult-to-sequence molluscs and arthropods, including millimeter-sized organisms. By highlighting collections as valuable sample resources and facilitating genome assembly of tiny and challenging organisms, our study advances efforts to obtain reference genomes of all eukaryotes.

## Background

High-quality genomes provide a powerful basis for understanding phylogenetic relationships, discovering fundamental principles of evolutionary processes, applying genomic methods to characterize, monitor and preserve biodiversity, and ultimately revealing the genetic blueprint underlying the fascinating diversity of life on our planet. Therefore, generating high-quality genomes of eukaryotic species has become a central goal in biological sciences [1]. Advances in short-read sequencing technology (with Illumina as the most prominent platform) enabled sequencing the genomes of a few thousand eukaryotes to date [2–5]. However, because eukaryotic genomes are often large and rich in repetitive DNA sequences, genome assembly from short reads ranging from 100 to 300 bp in size results in fragmented and incomplete assemblies [2,3,5], posing many limitations to downstream analyses. To generate highly contiguous genomes, the field has shifted to adopting long-read sequencing platforms from PacBio or Oxford Nanopore Technologies that can sequence DNA fragments with sizes of many kilobase pairs (kb) at once. Such long reads span most genomic repeats and do not suffer from the sequencing biases of short-read platforms in regions with very high or low GC content. Thus, long reads result in highly contiguous and complete genome assemblies, culminating in telomere-to-telomere assemblies [6,7], and consequently enable complete genome annotations and comprehensive analyses [8–16].

A key limitation of long-read sequencing is the availability of high molecular weight DNA, ideally with fragment sizes of 50 kb or more. To obtain samples delivering such DNA, the best practice is to acquire fresh samples (which may require sacrificing an individual), flash-freeze in liquid nitrogen, and preserve samples permanently at -80°C until DNA is extracted (https://www.earthbiogenome.org/sample-collection-processing-standards). Such protocols are not practical or not possible for (i) rare or endangered species, where sacrificing even a single living individual is not permitted, (ii) species which are difficult to sample in the field (e.g. cetaceans), or (iii) situations where liquid nitrogen and freezer capacity is not practicable (e.g. in remote areas). Therefore, sample availability is a key challenge for biodiversity genomics [17].

An alternative to get access to valuable or rare species that comprise Earth’s biodiversity are samples that are available in museums and other research collections that house millions of specimens worldwide, including samples from extinct species [18]. As one example demonstrating the value of such collections for biodiversity genomics, several hundred bird genomes have been generated from dry samples stored in museum collections [3]. However, since DNA of dry samples often exhibits various degrees of degradation, short-read sequencing was the only feasible technology, resulting in fragmented bird assemblies with an average contiguity of 43 kb (measured as contig N50 values, which state that 50% of the assembly consists of contiguous DNA segments – called contigs – of at least that size). Nevertheless, this and other studies using dry museum samples and short-read sequencing approaches, including marker-based sequencing and genome skimming, provided valuable insights into taxonomy, phylogenomics and conservation genomics [19–21].

In addition to dry material, collections worldwide also contain many millions of samples preserved in ethanol. In comparison to the logistical challenges associated with bringing liquid nitrogen to field trips and transporting flash-frozen samples without breaking the cold chain, preserving and transporting collected samples in ethanol is a notably simpler task. Since kilobase-sized DNA can be preserved in such samples [22,23], we explored whether ethanol-preserved samples are also suitable for long-read sequencing. We reasoned that even if DNA fragment sizes are substantially shorter than 50 kb, successfully sequencing reads of a few kilobases in size increases read length by at least an order of magnitude compared to short-read sequencing approaches, which in turn will improve assembly contiguity. In particular, we focused on the PacBio high-fidelity (HiFi) read protocol that instead of generating error-prone reads from “as long as possible” DNA fragments, sequences medium-sized fragments (10-15 kb) but with a high base accuracy of 99.8% [24]. HiFi sequencing enables assemblies that are both more contiguous and have a higher base accuracy than assemblies obtained with longer but more error-prone reads [7,16,24,25], making it a promising technology to apply to ethanol-preserved samples.

In this study, we explored the utility of ethanol-preserved samples from collections for HiFi sequencing. Although we encountered DNA degradation and sample contamination as expected problems in some samples, we also successfully demonstrate that HiFi reads can be obtained from ethanol-preserved samples containing kilobase-sized DNA, either using amplification-free protocols or by using a modified amplification-based protocol that effectively addresses issues associated with HiFi sequencing and PCR bias. Using this modified protocol, we generate a high-quality assembly of the 3.1 Gb genome of the maned sloth *Bradypus torquatus*, demonstrating that the previous genome size limit of 500 Mb can be substantially extended. Beyond collection samples, we further show that our modified protocol improves the contiguity of assemblies of species belonging to other phyla such as Mollusca (Gastropoda, Bivalvia) and Arthropoda (Collembola), where amplification is often required for long-read sequencing. The efficacy of this protocol facilitates genome assembly of challenging taxa and suggests that collections can serve as valuable sample sources for long-read sequencing.

## Results

### HiFi sequencing of ethanol-preserved samples with an amplification-free protocol

To investigate the effectiveness of PacBio HiFi sequencing from ethanol-preserved collection samples, we focused on vertebrates and used samples of four mammals (three-toed jerboa *Dipus sagitta*, pen-tailed treeshrew *Ptilocercus lowii*, long-eared flying mouse *Idiurus macrotis*, maned sloth *Bradypus torquatus*), two squamates (European blind snake *Xerotyphlops vermicularis*, slow worm *Anguis fragilis)* and two fishes (the catfish species *Cathorops nuchalis* and *Cathorops wayuu*), all lacking a genome assembly (Table 1, Supplementary Table 1). All samples were collected in the field and preserved in technical or 96% ethanol. Apart from the maned sloth and the catfishes, all samples were kept at room temperature. The samples of the maned sloth and catfish were kept most of the time in a freezer at -20°C; however, in contrast to flash-frozen samples, freezing did not occur immediately after sampling and they were kept at room temperature for extended periods of time, including during transportation.

**Table 1:**
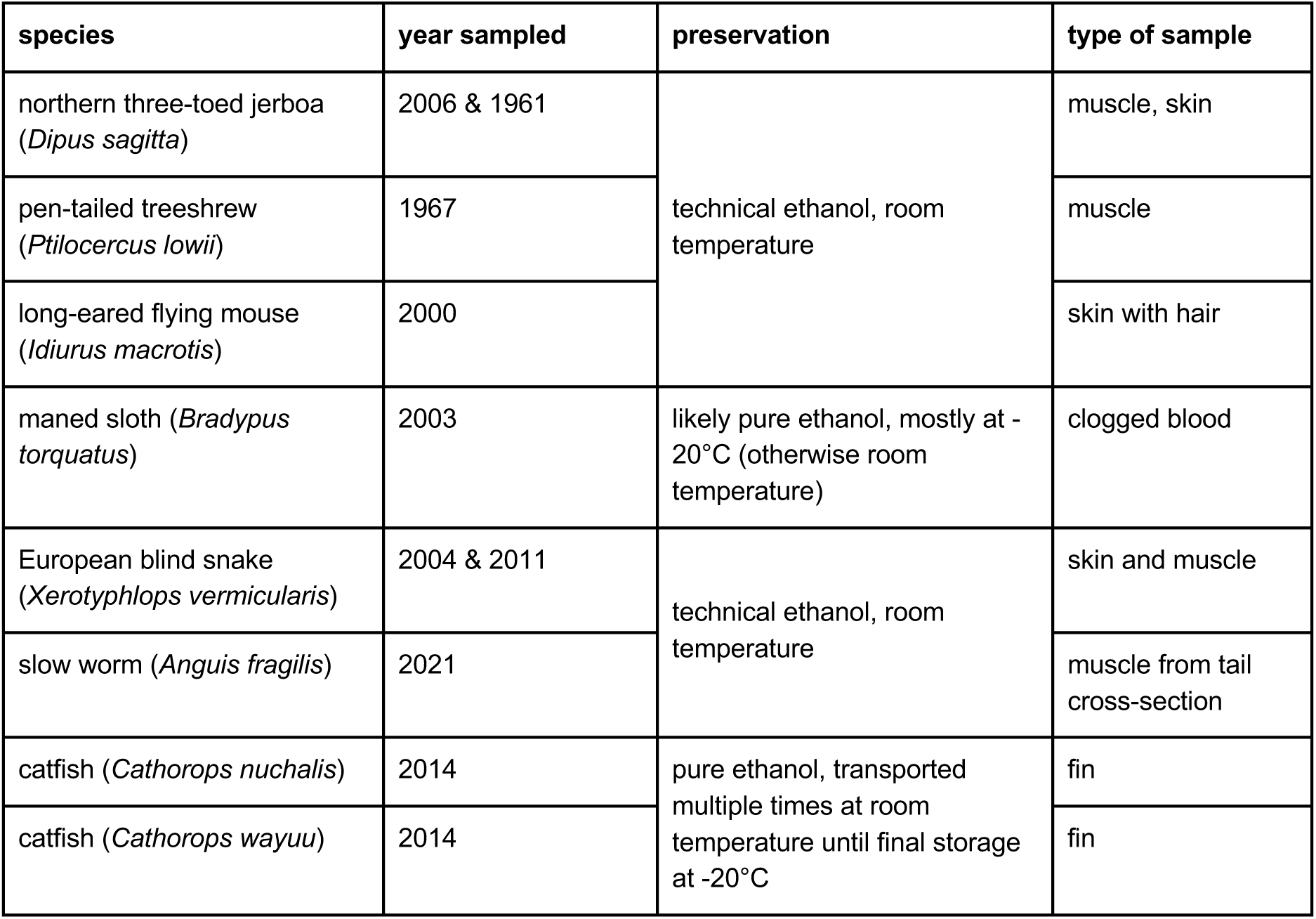
Overview of the species and samples.

| species | year sampled | preservation | type of sample |
| --- | --- | --- | --- |
| northern three-toed jerboa<br>(*Dipus sagitta*) | 2006 & 1961 | technical ethanol, room temperature | muscle, skin |
| pen-tailed treeshrew<br>(*Ptilocercus lowii*) | 1967 |  | muscle |
| long-eared flying mouse<br>(*Idiurus macrotis*) | 2000 |  | skin with hair |
| maned sloth (*Bradypus torquatus*) | 2003 | likely pure ethanol, mostly at -20°C (otherwise room temperature) | clogged blood |
| European blind snake<br>(*Xerotyphlops vermicularis*) | 2004 & 2011 | technical ethanol, room temperature | skin and muscle |
| slow worm (*Anguis fragilis*) | 2021 |  | muscle from tail cross-section |
| catfish (*Cathorops nuchalis*) | 2014 | pure ethanol, transported multiple times at room temperature until final storage at -20°C | fin |
| catfish (*Cathorops wayuu*) | 2014 |  | fin |

We used a modified Circulomics Nanobind disk and a phenol/chloroform based protocol for the extraction of genomic DNA (Methods). For *Dipus sagitta*, *Ptilocercus lowii*, and *Xerotyphlops vermicularis*, we did not obtain a sufficient amount of DNA (< 400 ng) and/or DNA fragments were shorter than 0.18 kb (Supplementary Table 1), showing that DNA is too degraded to proceed with library preparation. For four species (*Anguis fragilis*, *Idiurus macrotis*, *Cathorops nuchalis*, and *Cathorops wayuu*), the amount of DNA and the DNA fragment sizes were sufficient to prepare an amplification-free PacBio low input library [26] (Supplementary Table 1). We sequenced all libraries on a PacBio Sequel IIe system, disabling on-board calling of HiFi reads and instead applying the computationally expensive DeepConsensus method [27] to maximize HiFi read yield and length. For *Bradypus torquatus*, we did not obtain enough DNA and therefore proceeded with a PacBio ultra-low input library (see below).

For the two catfish species, *Cathorops nuchalis* and *Cathorops wayuu*, we sequenced two SMRT cells each and obtained HiFi reads with an average length of 8,832 and 8,783 bp, respectively, providing a total of 43.8 and 41.2 Gb, which corresponds to a coverages of ∼17X and ∼16.5X (Supplementary Table 1). Using HiFiasm with different parameters [28], we were able to obtain a contig assembly for both species with a total length of 2.6 and 2.59 Gb and a contig N50 value of 3.2 and 2.1 Mb (Supplementary Table 2). To assess gene completeness, we used compleasm [29] with the set of 3,640 ray-finned fish (Actinopterygii) near-universally conserved genes (ODB10) [30], which showed that 96.65% of these genes are fully present in the primary assembly of *C. nuchalis* and 95.6% in that of *C. wayuu*. Although additional HiFi data would be needed to improve contiguity and HiC data would be required to scaffold the contigs into chromosome-level scaffolds, our catfish samples exemplify that an adequate genome assembly can be obtained from 10-year-old, ethanol-preserved tissues.

In contrast to the catfish, we obtained very low sequencing yields for *Idiurus macrotis* and *Anguis fragilis*, with only 0.3 Gb and 0.04 Gb of HiFi data (Supplementary Table 1). Quality metrics showed that the polymerase N50 raw read lengths were very short and the local base rates were low. For example, while the library from *Anguis* met the requirements for PacBio sequencing with a mean fragment length of 12.2 kb, both the local base rate of 1.64 (expected ∼2.8) and the polymerase N50 raw read length of 32.3 kb (expected at least 200 kb) are very low and insufficient to produce HiFi reads of most DNA fragments in the library. This indicates that factors such as DNA damage, metabolites bound to the DNA, or contaminants precipitated with the DNA inhibit the polymerase, highlighting sequencing challenges for ethanol-preserved samples stored at room temperature.

### HiFi sequencing with the amplification-based ultra-low input protocol

We reasoned that a PCR-based amplification step prior to library preparation could render the *Idiurus macrotis* and *Anguis fragilis* samples amenable to sequencing, as this procedure should yield intact DNA devoid of potential polymerase-inhibiting metabolites. To this end, we applied the PacBio ultra-low input library protocol [31] to the samples of *Idiurus macrotis* and *Anguis fragilis*. Although this protocol was originally designed for small specimens providing very limited DNA amounts [32] and is recommended only for genome sizes of up to 500 Mb, the protocol includes a PCR amplification step using two different undisclosed polymerases targeting DNA with average and high GC contents, respectively. For simplicity, we refer to these polymerases as “A” and “B” in the following to distinguish them from a third polymerase “C” that we also investigate as described below. We also generated an ultra-low input library for the *Bradypus torquatus* sample that did not contain enough DNA for the low input protocol. Indeed, for *Idiurus macrotis* and *Anguis fragilis*, sequencing another SMRT cell each produced 10 and 19.6 Gb in HiFi reads with an average HiFi read length of 4,854 bp and 7,552 bp. The first SMRT cell for *Bradypus torquatus* yielded 29.9 Gb in HiFi reads with an average HiFi read length of 10,850 bp (Supplementary Table 1).

For *Idiurus macrotis* and *Anguis fragilis*, we investigated whether a DNA repair step applied to the DNA extract before preparing the ultra-low library would increase HiFi read length and yield (Methods). In contrast to the previous sequencing results, adding the DNA repair step produced shorter HiFi reads (average read length 4,270 vs. 4,854 bp for *Idiurus macrotis* and 5,609 vs. 7,552 bp for *Anguis fragilis*) and a lower yield (6.4 vs. 10 Gb for *Idiurus macrotis* and 12.6 vs. 19.6 Gb for *Anguis fragilis*), suggesting that the DNA repair process is not advantageous for these samples (Supplementary Table 1).

Next, we investigated whether the sequenced DNA was contaminated with bacteria, fungi or other microorganisms. While little contamination was found in the *Bradypus torquatus* sample (∼200 kb mostly assigned to plants), the *Anguis fragilis* data had higher levels of contamination (∼200 Mb assigned to various bacterial groups), and the vast majority of the sequencing data obtained from the *Idiurus macrotis* sample were contamination (∼75 Mb assigned to various groups of bacteria) (Supplementary Figures 1, 2). High levels of contamination (71-90% of sequenced reads) were also detected for three additional ethanol-preserved samples, where we directly applied the ultra-low input protocol: Russian desman (*Desmana moschata*) sampled in 1947, Hazel dormouse (*Muscardinus avellanarius*) sampled in 2016, and a *Anguis fragilis* sample from 1878 (Supplementary Tables 1, 3). Together, while sample contamination with bacteria, protists and bacterial viruses or cross-contamination with human DNA is another challenge related to samples obtained from collections [33–35], our tests also show that amplifying DNA in the ultra-low input protocol prior to library preparation can enable PacBio HiFi sequencing of samples where the amplification-free low input library protocol failed.

### PCR bias in the current protocol prevents high-quality assemblies of larger genomes

To investigate the feasibility of using the ultra-low input protocol to obtain a high-quality assembly of a genome that substantially exceeds the recommended size limit of 500 Mb, we focused on the maned sloth that has an estimated genome size exceeding 3 Gb and showed a low level of contamination. To obtain sufficient read coverage for genome assembly, we generated two additional libraries using the PacBio ultra-low protocol and sequenced four additional SMRT cells. In total, all five SMRT cells provided 140.2 Gb of HiFi reads, a total coverage of ∼45X, with an average read length of 10.6 kb. However, using this data, we only obtained an assembly with a contig N50 of 405 kb (Figure 1, brown dashed line), which is unexpectedly low as similar read coverages typically yield mammalian assemblies with contig N50 values exceeding several megabases. Using compleasm [29] with the set of 11,366 near-universally conserved eutheria genes (ODB10) showed that only 85.3% of these genes are fully present in our assembly. Similarly, using TOGA [36] to determine how many of the 18,430 ancestral placental mammal coding genes have an intact reading frame, revealed that only 68% of the ancestral genes are intact. Together, this indicates not only a low assembly contiguity but also a high level of incompleteness.

**Figure 1:**
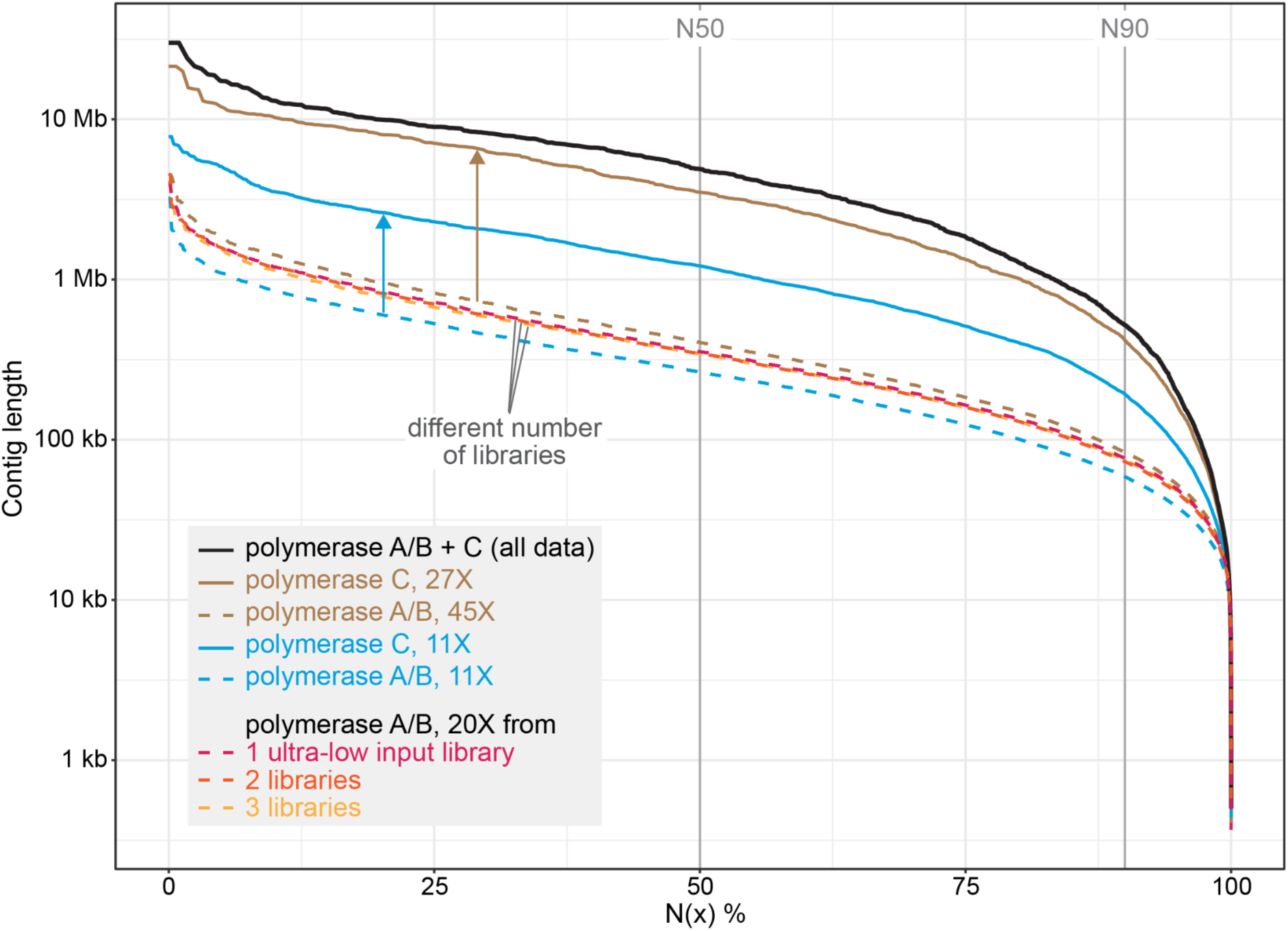
Contiguity of *B. torquatus* assemblies generated with data from ultra-low input libraries prepared with polymerases A/B and/or C at different coverages. Assembly contiguity visualized as N(x) graphs that show contig sizes on the Y-axis, for which x percent of the assembly consists of contigs of at least that size. The N50 and N90 values are shown as vertical grey lines and indicate contig sizes for which 50% and 90% of the assembly consists of contigs of at least that size, respectively. Assemblies involving polymerase C read data are shown as solid lines, assemblies generated from polymerase A/B data are shown as dashed lines. Colors refer to different comparisons discussed in the text and summarized in the inset.

To further investigate the reasons for the poor quality of this assembly, we aligned our *Bradypus torquatus* HiFi reads against the high-quality genome of a related sloth species, *Choloepus didactylus* [11]. Despite both species being separated for 30 My [37], we observed that 84.3% of the *Choloepus didactylus* genome was covered with *B. torquatus* HiFi reads at an average coverage of 38X. Inspecting the mapped reads in a genome browser revealed larger genomic regions, often spanning many kilobases, that completely lack any mapped reads (Figure 2). Since several of these regions contain highly-conserved genes, we reasoned that these read dropouts are probably not caused by high divergence between the sloth species. Instead, it is likely that despite relying on two polymerases, the PacBio ultra-low protocol has PCR bias on larger genomes, resulting in genomic regions that lack any reads.

**Figure 2:**
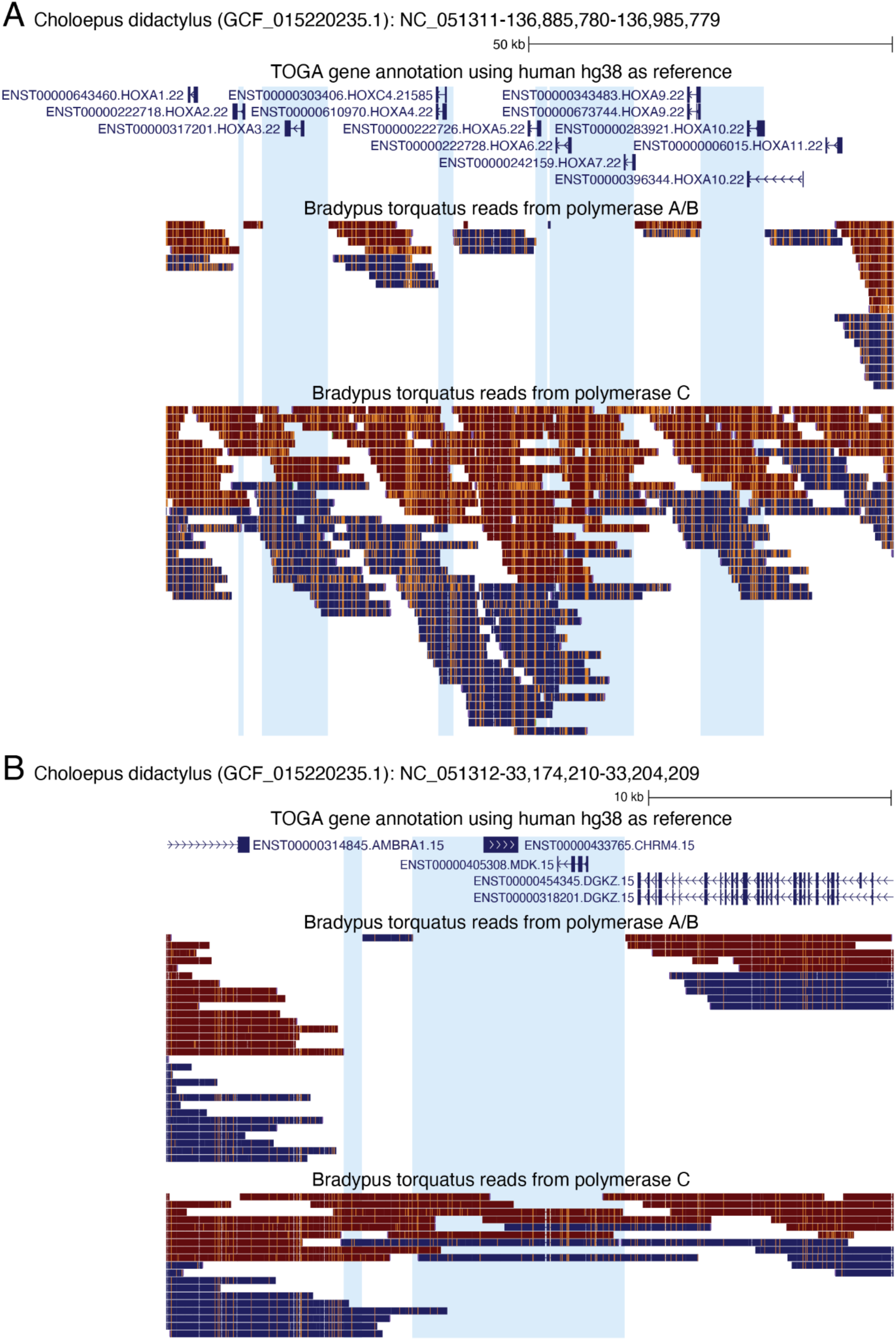
PCR bias in reads produced with polymerase A/B. UCSC genome browser screenshots of the *Choloepus didactylus* assembly, together with the TOGA gene annotation and mapped HiFi reads of *B. torquatus* produced either with polymerase A/B or polymerase C. The TOGA gene annotation is shown in blue with boxes representing coding exons, connecting horizontal lines representing introns, and arrowheads indicating the direction of transcription (+ or - strand). Mapped HiFi reads are shown below as boxes with orange tickmarks representing insertions in the *B. torquatus* reads relative to the *C. didactylus* assembly. Reads in blue and red align to the + and - strand, respectively. (A) In the *HOXA* gene cluster, several regions, often covering parts or entire *HOX* genes, lack any reads produced with polymerase A/B (highlighted in blue). In contrast, these regions have a coverage of HiFi reads produced with polymerase C, which is sufficient for assembly. (B) While reads produced with polymerase A/B do not cover the *CHRM4* and *MDK* genes, polymerase C reads cover the entire locus.

### A different polymerase alleviates PCR bias and enables highly-complete assemblies of larger genomes

To alleviate PCR bias, we adapted the ultra-low input protocol and used a different polymerase, KOD Xtreme™ Hot Start DNA Polymerase (Merck). According to the specification sheet, this polymerase amplifies DNA fragments up to 24 kb at high fidelity, including templates with up to 90% GC content, which could help to overcome the underrepresentation of very low or high GC regions of the PacBio ultra-low input protocol. For simplicity, we refer to this polymerase as “C” in the following. Using a single library, we sequenced another three SMRT cells for the maned sloth, providing 91.7 Gb (corresponding to an additional 27X coverage) of reads with an average length of 10.2 kb.

Performing genome assembly using all HiFi reads obtained with polymerase A/B and C, produced a 3.13 Gb assembly with a contig N50 of 4.88 Mb (Figure 1, black line, Supplementary Table 4), which is 12 times higher than the previous assembly generated from reads obtained with polymerase A/B. Gene completeness estimated with compleasm improved from 85.3% to 96.4% and the percentage of intact ancestral placental mammal genes inferred with TOGA increased from 68 to 88.6%. Furthermore, mapping the polymerase C HiFi reads to the *C. didactylus* assembly covered the regions that completely lacked any read before (Figure 2). Consistent with a higher PCR bias for polymerase A/B, we found that the normalized read coverage in exonic and repeat regions is biased towards a lower coverage for the polymerase A/B data compared to polymerase C to data (Supplementary Figure 3). This confirms that previous read dropouts were not caused by sequence divergence between both sloth species or selective degradation of certain genomic regions in our sample, but by PCR bias associated with the polymerases in the PacBio ultra-low input protocol.

To directly compare the effect of polymerase A/B vs. C, taking differences in read coverage from the individual SMRT cells out of the equation, we downsampled our data to equal coverage and performed a number of tests (Supplementary Table 4). Since DNA fragments generated by polymerase A and B are pooled during the library preparation, we cannot investigate the effect of those two polymerases individually. Using an equal, downsampled coverage of ∼11X, we found that the assembly produced from only polymerase C reads outperformed the assembly produced from only polymerase A/B reads by exhibiting a substantially higher contiguity (contig N50 1.22 Mb vs. 264 kb) and gene completeness (89.3% vs. 77.0% completely detected genes) (Figure 1, light blue lines). Remarkably, an assembly obtained from the complete polymerase C read data is substantially better than an assembly obtained from the complete polymerase A/B read data (contig N50 3.5 Mb vs. 405 kb, 96.4% vs. 85.3% completely detected genes) (Figure 1, brown lines), despite the polymerase C data having a substantially lower coverage (27X vs. 45X for polymerase A/B).

We next investigated how the number of libraries produced with polymerase A/B influences assembly, as additional libraries may increase complexity and reduce bias. However, sampling an equal coverage of ∼20X from either one, two or three libraries results in very similar assemblies in terms of contiguity and gene completeness (Figure 1, red/orange/yellow lines; Supplementary Table 4), indicating that inherent bias of polymerase A/B hampers assembly quality that cannot be overcome by producing several libraries.

Together, these tests show that *B. torquatus* assemblies generated with polymerase C reads are substantially better. To our knowledge, we provide the first high-quality contig assembly of a 3.1 GB genome that was produced using an adapted ultra-low input protocol combining polymerase A/B and C.

### Chromosome-level assembly of the maned sloth

To obtain a final scaffolded assembly of *Bradypus torquatus*, we used the Arima HiC protocol, which is applicable to ethanol-preserved samples [23,38], to generate 97.5 Gb in long-range read pair data. Using the automated scaffolding software yahs [39] and manual curation, our contig assembly could be scaffolded into chromosome-level scaffolds (Figure 3A). This final assembly consists of 2,915 scaffolds and 5,022 contigs. The scaffold N50 and N90 values are 157 Mb and 61.3 Mb, respectively (Figure 3B). The contig N50 and N90 values are 4.75 Mb and 519 kb, respectively. Using Merqury [40] with the HiFi reads, we estimate a high base accuracy (QV=46.7), which represents an upper bound as the HiFi reads were also used for assembly. The assembly has a compleasm gene completeness score of 97.3% based on the eutheria ODB10 database with n=11,366 genes (Supplementary Table 4) and contains 90.72% of ancestral placental mammal genes.

**Figure 3:**
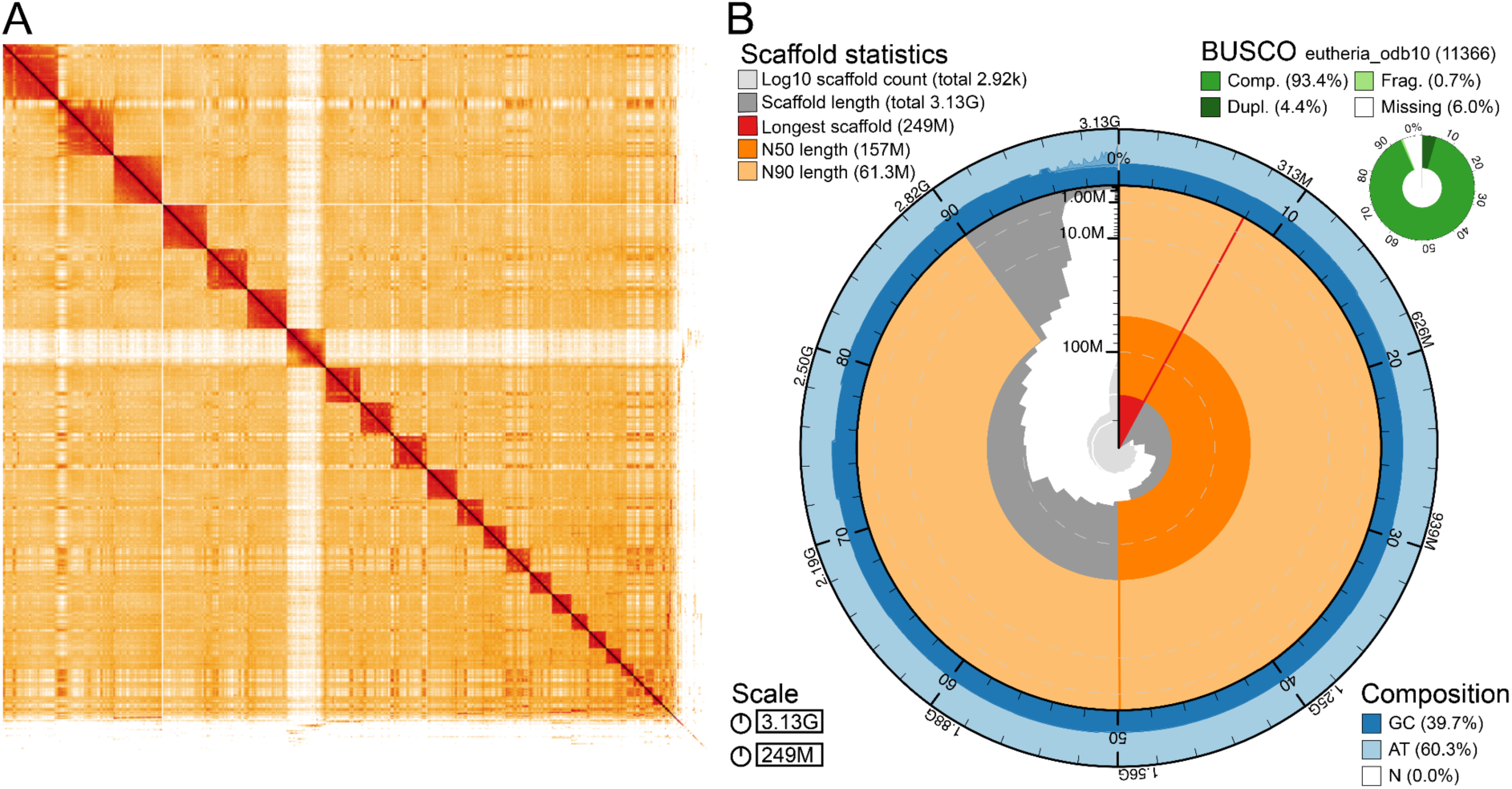
Chromosome-scale assembly of *Bradypus torquatus*. (A) HiC interaction map after automated scaffolding by yahs and manual curation. The HiC map shows interactions in 3-dimensional space between two regions of the genome. Darker colors indicate a higher number of interactions. The region of low interaction between scaffold 7 and all other scaffolds indicates this scaffold is the X chromosome, which was confirmed as this scaffold aligns to the human X chromosome. (B) Snail plot showing lengths of all scaffolds, together with the longest scaffold (red), and the N50 (dark orange) and N90 length (light orange). The outer ring shows the GC content of the genome.

In comparison to existing genome assemblies of xenarthran species, our final assembly clearly outperforms the short-read based assembly of the sloth *Choloepus hoffmanni* in terms of contiguity and the number of intact ancestral placental mammal genes (Figure 4). Although other long-read based xenarthran assemblies, which were most likely generated from flash-frozen samples obtained from zoos and captive colonies, have even higher contiguities, our *Bradypus torquatus* assembly is a valuable addition for xenarthran and, more generally, mammalian comparative genomics.

**Figure 4:**
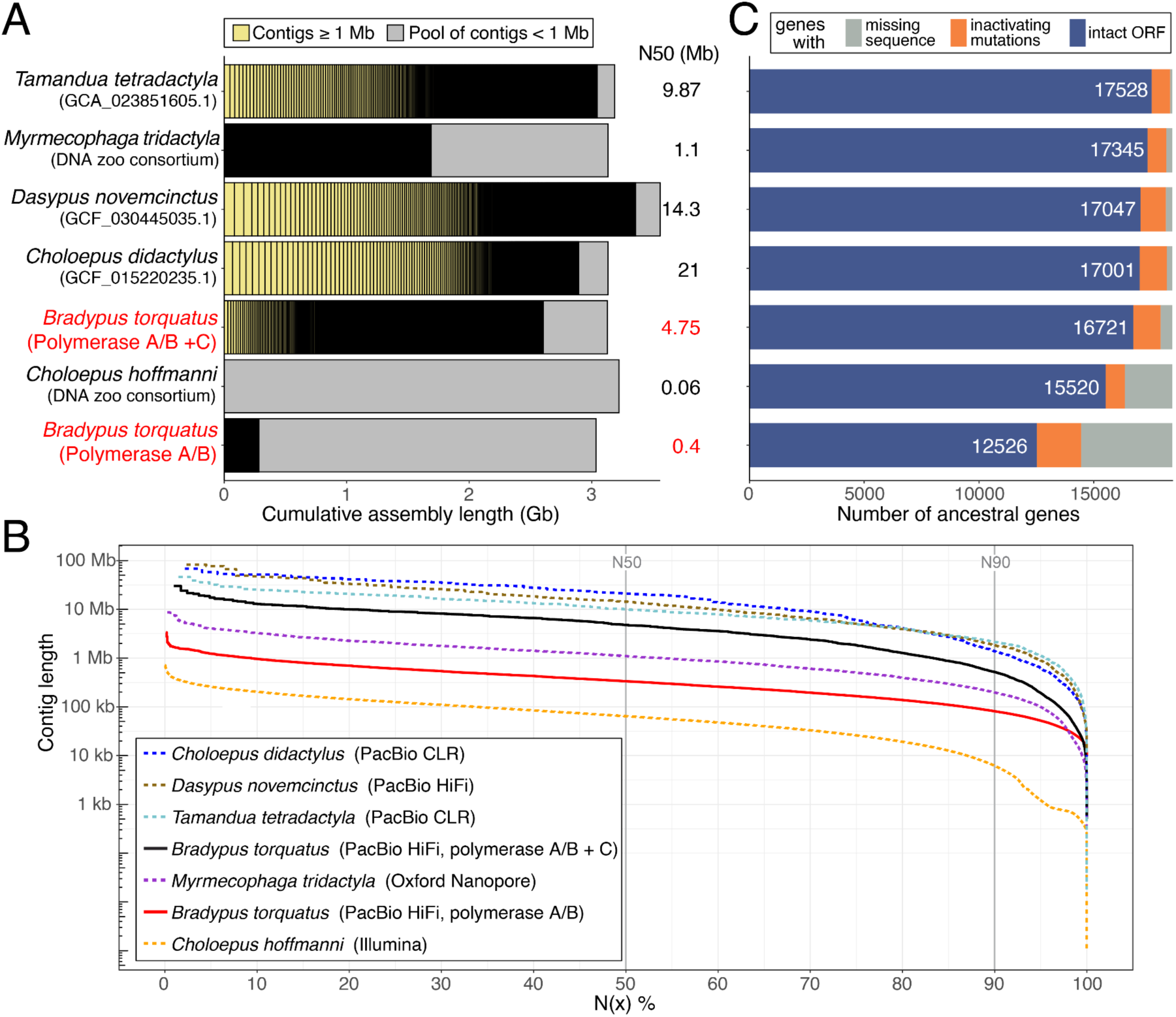
Comparison of xenarthran genome assemblies. (A) Visualization of contig sizes of available xenarthran genome assemblies. Each bar represents the total assembly size. Contigs shorter than 1 Mb are not visualized individually, but shown as the grey portion of each bar. The final *B. torquatus* assembly and its preliminary assembly generated only from polymerase A/B reads are in red font. Assembly source or accession is listed in this panel, the sequencing technology used is listed in the inset in panel B. (B) Visualization of assembly contiguity as an N(x) graph, showing contig sizes on the Y-axis, for which x percent of the assembly consists of contigs of at least that size. Assembly order in the legend (inset) is sorted by contig N50 value. (C) TOGA classification of 18,430 ancestral placental mammal genes showing the number of genes that have an intact reading frame (blue bar, number is given in white font), inactivating mutations (e.g. frameshifts, stop codon, splice site mutations or exon deletions; orange bar), or missing coding sequence parts often caused by assembly gaps or fragmentation (gray bar). Assemblies are sorted by the number of intact genes.

### Polymerase C improves assemblies for various species

We next explored whether polymerase C can also help to improve assemblies of other species, using samples not obtained from collections. To provide a fair comparison, we randomly downsampled the larger data set to obtain an equal coverage of HiFi reads generated with polymerases A/B and C. To compare these polymerases for another mammal, we used the human HG002 sample and generated assemblies for both human haplotypes. Using an equal coverage of 23.5X, the polymerase A/B read data produced a 2.96 Gb assembly for haplotype 1 with a contig N50 value of 642 kb, whereas the polymerase C data generated a 3.03 Gb assembly with a substantially higher contig N50 value of 2.8 Mb, a 4.4 fold increase in contiguity. Consistently, gene completeness assessed with compleasm (mammalia_odb10) increased substantially from 81.2 to 98.6%. Similar results were obtained for the haplotype 2 assembly, where the polymerase A/B read data produced a 2.9 Gb assembly with a contig N50 value of 558.8 kb and a gene completeness of 77.3%, whereas the polymerase C read data produced a 3 Gb assembly with a contig N50 value of 2 Mb and a gene completeness of 97.8%.

Since PCR amplification may produce chimeric reads [41], we used available non-amplified human HiFi reads produced from the HG002 sample as a baseline to compare the amount of chimeric HiFi reads generated by polymerase A/B and C. We mapped reads to the HG002 assembly [6] and computed the number of reads with supplementary alignments, which indicate chimers. We found that the fraction of chimeric alignments is very low (≤0.81%) across all three libraries, with polymerase C reads having the lowest fraction (Supplementary Figure 4A). We next included available HG002 read data obtained by Multiple Displacement Amplification (MDA) in this comparison. Consistent with previous observations [41,42], the majority of MDA alignments (69.3%) are chimers, which is further supported by the observation that the primary alignment lengths are much shorter than the MDA reads (Supplementary Figure 4). We therefore conclude that long range PCR amplification used in the original and modified ultra-low input protocol does not create more chimeric reads than non-amplified libraries and orders of magnitude fewer chimeric reads than MDA libraries.

Next, we explored the application of polymerase C to three non-vertebrate taxa covering two additional phyla, Mollusca (two taxonomic classes: Gastropoda and Bivalvia) and Arthropoda (Collembola), using taxa where genome sequencing efforts often rely on the amplification-based protocols because of low sequencing performance with low input protocol or very small DNA amounts.

For the sacoglossan gastropod *Elysia timida* (Mollusca), previous sequencing libraries created with the low input protocol resulted in very poor sequencing performance. Therefore, we applied the ultra-low input protocols, and compared two SMRT cells produced with polymerase A/B, providing 16.6 and 20.8 Gb yield in reads with an N50 length of 6.5 and 5.8 kb, to one SMRT cell produced with polymerase C, providing 23 Gb yield in reads with an N50 length of 7 kb (Supplementary Table 5). After subsampling to equal read coverage of 26.4X, polymerase A/B and C read data generated assemblies with similar contig N50 values of 347.1 kb for polymerase A/B and 331 kb for polymerase C (Figure 5, Supplementary Table 5). Using all polymerase A/B read data with a coverage of 42.5X increased the contig N50 value to 472.6 kb. Importantly, adding the 23 Gb of polymerase C reads, increased the contig N50 value 1.4 fold to 675.8 kb (Figure 5). While the gene completeness (metazoa_odb10) of 97.7 and 97.8% is similar between these assemblies, polymerase C data helped to improve assembly contiguity for this mollusc.

**Figure 5:**
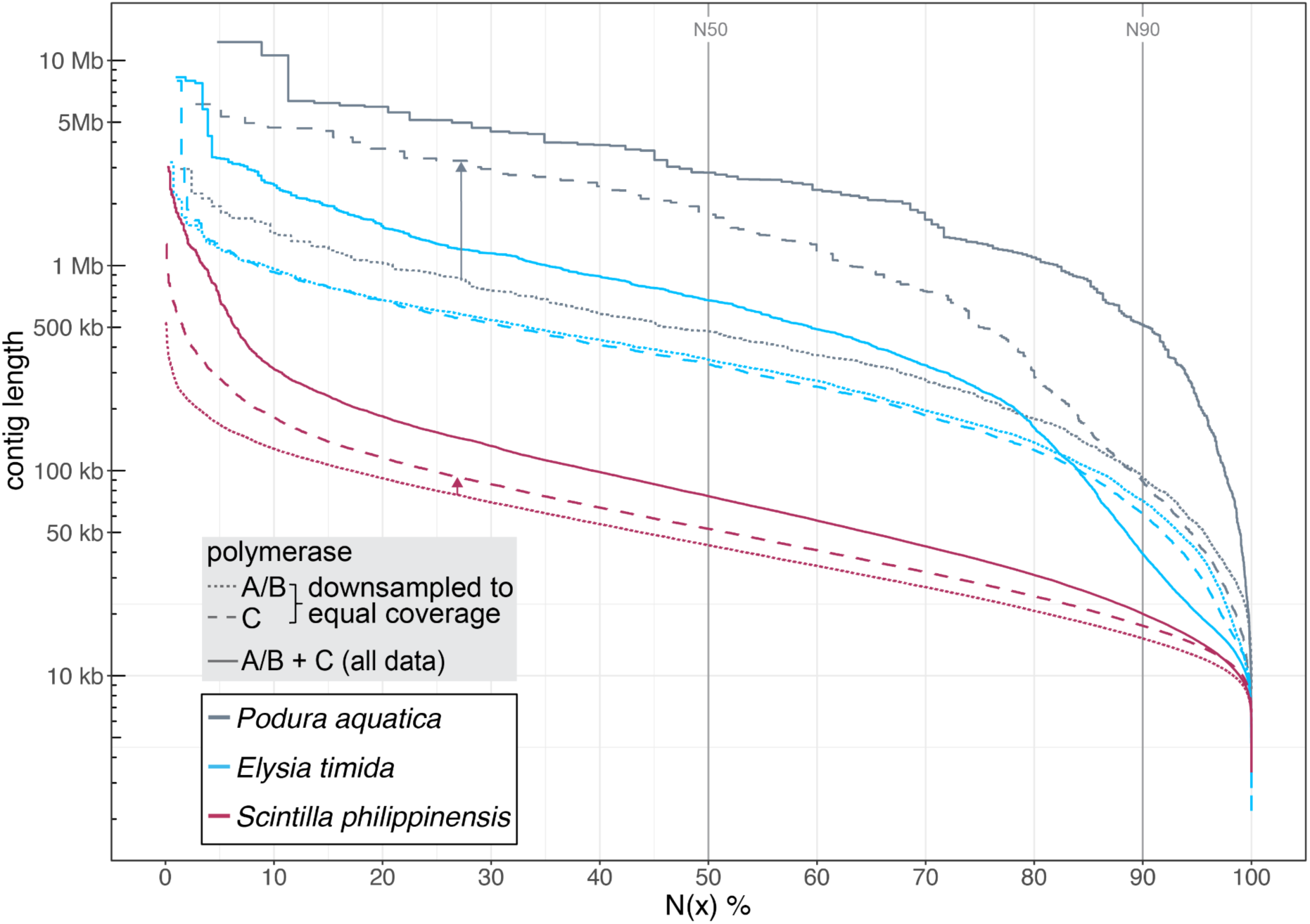
Impact of polymerase C on assemblies of mollusc and collembola species. Assembly contiguity visualized as N(x) graphs that show contig sizes on the Y-axis, for which x percent of the assembly consists of contigs of at least that size (N50 and N90 values are indicated). Assemblies are generated with an equal (downsampled) coverage of reads from polymerase A/B (dotted lines) and C (dashed lines). Assemblies generated with all data are shown as solid lines. Colors refer to different species.

To understand why polymerase C alone does not result in a more contiguous assembly, we mapped both polymerase A/B and C reads to the *Elysia timida* assembly with highest contiguity, generated from all read data. This showed that both polymerases A/B and C exhibit bias; however, bias of one polymerase can be compensated by reads of the other (Supplementary Figure 5), indicating that these polymerases may have taxon-specific differences.

For the marine bivalve *Scintilla philippinensis* with an estimated genome size of 1.3 Gb, we compared assemblies produced from 22.3 Gb of reads obtained from ultra-low input libraries using polymerase A/B or C, which corresponds to a coverage of 17.1X. While the polymerase A/B reads produced a 1.77 Gb assembly with a contig N50 value of 43.3 kb, the polymerase C read data produced a 1.88 Gb assembly with a 1.2 fold increased contig N50 value of 52.1 kb. Gene completeness (metazoa_odb10) improved slightly from 89.1% (polymerase A/B) to 89.9% (polymerase C). Combining all polymerase A/B and C read data (coverage of 36.1X) produced a 1.86 Gb assembly with an even higher contig N50 value of 75.1 kb (Figure 5, Supplementary Table 5) and a higher gene completeness of 93.5%. Mapping polymerase A/B and C reads to the assembly generated with all data also revealed regions that were covered only by reads from one polymerase (Supplementary Figure 6A,B). While polymerase C reads improved assembly contiguity of both mollusc species, the resulting assemblies have a comparatively low contiguity, highlighting the challenges of sequencing molluscan DNA.

We next tested our adjusted protocol on a species having a very small body size, where amplification of the limited amount of genomic DNA is required for long-read sequencing and genome assembly [32]. We used an ethanol-preserved, whole single specimen of the springtail *Podura aquatica* (Arthropoda: Collembola), which has a body size of only 1.5 mm and an expected genome size of 200-300 Mb. The polymerase A/B run yielded 13.9 Gb of HiFi reads with an N50 length of 9.6 kb. The polymerase C run yielded 21.7 Gb of reads but with a lower N50 read length of 5.7 kb, which is likely explained by sequencing DNA one year after the initial extraction (the entire specimen was used for the initial DNA extraction). Strikingly, at an estimated coverage of ∼50X, the polymerase A/B read data produced a 278.5 Mb assembly with a contig N50 value of only 919 kb, whereas the polymerase C data generated a 269.3 Mb assembly with a contig N50 value of 2.77 Mb (Figure 5, Supplementary Table 5). This represents a 3 fold increase in contiguity, despite the polymerase C reads being substantially shorter. Gene completeness (arthropoda_odb10) increased slightly from 92.8% for polymerase A/B assembly to 93.4% for polymerase C assembly. Combining all polymerase A/B and C read data resulted in a 284.7 Mb assembly with an even higher contig N50 value of 5.74 Mb and the same gene completeness of 93.4%. Similar to *Elysia* and *Scintilla*, aligning reads to the most contiguous assembly showed complementary coverage dropouts (Supplementary Figure 6C,D).

Together, these tests confirm that polymerase C improves the assembly contiguity and sometimes gene completeness for a broad range of species, including species that rely on amplification-based library preparation protocols because their small size does not provide enough DNA from a single individual or because naturally-occurring metabolites presumably inhibit the polymerase during sequencing.

## Discussion

Our investigation into utilizing collection samples for long-read sequencing confirms that ethanol-preserved samples can contain kilobase-sized DNA, long enough for long-read sequencing [22,23]. For the two catfish species, we found that amplification-free protocols generated sequencing data sufficient to generate assemblies with contig N50 values surpassing 2 Mb. Application of amplification-free protocols is recommended whenever feasible, as they will not suffer from PCR bias. Our other tests indicate that mammal or reptile samples may necessitate amplification-based protocols. It remains to be investigated for which taxonomic groups amplification-free protocols are generally successful. We demonstrate that PCR bias associated with the amplification-based PacBio ultra-low input protocol can be overcome or at least mitigated by employing an alternative polymerase. As a proof of concept, the contiguous 3.1 Gb genome assembly of *B. torquatus* shows that a modified amplification-based protocol can produce high-quality assemblies of gigabase-sized genomes.

Contamination caused by sample decomposition, human handlers, or commensal bacteria is expected for collection samples that have been stored under non-sterile conditions [33]. It is difficult to assess contamination prior to sequencing, and we find different levels of contamination in our samples, ranging from most of sequenced reads stemming from contaminants to almost no contamination. Analyzing a low coverage of sequencing reads for contamination before sequencing a sample to the coverage required for assembly could therefore be a cost-efficient strategy to select those samples that contain sufficiently low contamination levels. Furthermore, the resulting assemblies should be carefully screened for contamination using existing methods [43,44].

Consistent with previous observations [34], we find that sample age alone is not an accurate predictor of input DNA quality and sample suitability for sequencing. For example, while the *B. torquatus* sample was collected in 2003, several younger samples exhibited high degrees of DNA degradation (Supplementary Table 1). Hence, in addition to sample age, other factors such as storage temperature and conditions, storage medium, or tissue type likely influence DNA quality. From our experience, samples consistently stored at -20°C and preserved in 96% ethanol perform well, but a systematic assessment of larger sample numbers is needed to substantiate this.

Our study has a number of implications. First, the modified ultra-low input protocol improves genome assembly of small specimens, where amplification is a requirement to obtain enough DNA for sequencing. For example, the contiguity of the *Podura aquatica* genome increased to an N50 of 5.7 Mb, and thus substantially exceeds the minimum standards of 100 kb set by the Earth Biogenome Project for small species with limited DNA amounts [1]. The modified protocol will likely not only be beneficial for species with diminutive body sizes that represent a very large but mostly uncharacterized part of Earth’s biodiversity, but also in cases where only very limited amounts of material from non-lethal samplings (biopsies from human patients or bat wing punches) are available. Second, long-read sequencing remains a challenge for molluscs and other taxonomic groups, where satisfactory sequencing outputs often require amplification-based protocols. Although achieving highly contiguous assemblies with megabase contig N50 values remains challenging for these species, our investigations suggest that employing a combination of different polymerases can at least help to improve assembly contiguity. Third, while the PacBio ultra-low input protocol was previously limited to genome sizes of up to 500 Mb, the successful application of the modified protocol to *B. torquatus* with its 3.1 Gb genome extends its applicability to a broad range of species with larger genome sizes. Together, the improved efficiency of the modified ultra-low input protocol opens avenues for generating contiguous genomes across various species.

Our study raises the question of finding polymerases with minimal bias. While our tests with *B. torquatus* and human indicate that polymerase C shows satisfactory performance for mammals, we found that polymerase C also appears to exhibit bias for samples of molluscs and collembola, albeit a different bias compared to polymerase A/B (Supplementary Figures 5, 6). Anticipating that DNA amplification will constitute a key step in the genome sequencing procedure for numerous collection samples, challenging species, and species with diminutive body sizes, future investigations could focus on identifying the most appropriate polymerase or combination of polymerases that exhibit minimal bias for specific taxonomic groups.

Apart from the ultra-low input protocol, several new approaches have recently been developed to make small amounts of input DNA accessible for long read sequencing. This includes the above-mentioned MDA [41,42,45], adapter ligation via tagmentation [46,47], and Picogram input multimodal sequencing (PiMmS) [48]. We show here that the ultra-low input protocol produces very few chimeric reads in contrast to MDA. Furthermore, the ultra-low input protocol can generate average read lengths of ∼10 kb, which is similar to read lengths generated by PiMmS [48], but substantially longer than those generated with tagmentation based approaches (2.5-5 kb averages) [46,47]. Nevertheless, different methods likely have ideal application ranges that depend on the input sample, its quality and amount of DNA. Future research should therefore benchmark which library preparation method is optimal for which sample type.

## Conclusions

Our work suggests that collections can complement flash-frozen material as a sample source for biodiversity genomics, especially for species that are hard to sample because of rarity, protection status or other reasons. Thus, natural history collections as extensive archives of biodiversity can help to achieve the ambitious goal of generating reference genomes for all life on Earth.

## Material and Methods

### Sample sources

For *Bradypus torquatus*, we used a sample of ∼50 mg of clogged blood, preserved in ethanol. This sample was collected in 2003 under Brazilian license SISGEN number AF86294 and CITES number 138261. For *Idiurus macrotis*, we used ∼12 mg of skin with hair preserved in technical ethanol at room temperature. For the *Anguis fragilis*, we used ∼51 mg (for the one collected in 2021) and ∼3 mg (for the one collected in 1878) of muscle tissue from a tail cross-section. Both samples were preserved in technical ethanol at room temperature. For both *Cathorops* species, we used fin samples stored in ethanol in frozen collections at the Leibniz Institute for the Analysis of Biodiversity Change (LIB) Bonn. Originally, fin clips of specimens acquired from local fishermen were taken in 2014, immediately placed into ethanol, but subsequently transported multiple times at room temperature until final storage at -20°C. The exact time between catch and sampling is unknown but was likely a few hours. For *Desmana moschata*, we used ∼9 mg of muscle and skin tissue that was preserved in ethanol at room temperature. For *Muscardinus avellanarius*, we used ∼19 mg of foot tissue that was preserved in technical ethanol at room temperature. For *Dipus sagitta*, we used ∼12 mg (individual 95545) and ∼16 mg (individual 95541) of muscle tissue and ∼30 mg (individual 56492) of skin. All three samples were preserved in technical ethanol at room temperature. For *Ptilocercus lowii*, we used ∼8 mg of muscle tissue that was preserved in technical ethanol at room temperature. For *Xerotyphlops vermicularis*, we used ∼5 mg (individual collected in 2004) and ∼3 mg (individual collected in 2011) of skin and muscle tissue preserved in technical ethanol at room temperature. For *Elysia timida*, we used a whole specimen (∼1 cm body length) from our living culture, which we immediately homogenized for DNA extraction after euthanization. This sample was collected under license ESNC 205 issued by the Spanish “Dirección General de Biodiversidad, Bosques y Desertificación del Ministerio para la Transición Ecológica y el Reto Demográfico”. For *Scintilla philippinensis*, we used ∼20 mg of muscle tissue preserved in ethanol, collected in Johor Malaysia under a collaboration agreement between Senckenberg and Universiti Putra Malaysia. For *Podura aquatica*, we used a single whole specimen (∼1.5 mm body length) killed and immediately preserved in 96% ethanol. Two libraries were produced either with polymerase A/B or polymerase C (below), and while the polymerase A/B experiment was done within the month following DNA extraction, the polymerase C experiment was conducted one year after DNA extraction, using DNA preserved at -20°C in TE buffer. Supplementary Table 1 lists sample sources, accessions and additional details.

### DNA extraction

High molecular weight (HMW) gDNA was extracted from ethanol-preserved tissues of *Bradypus torquatus*, using a modified protocol version of the Circulomics Nanobind Tissue Big DNA kit, including the ethanol removing step described in ‘Guide and overview – Nanobind tissue kit’. We retrieved gDNA bound to the Nanobind disk as well as unbound gDNA in the precipitation solution. The gDNA bound to the Nanobind disk was eluted after several washing steps. The unbound gDNA in the precipitation solution was precipitated by centrifugation (18.000 xg for 30 min at 4°C). The resulting pellet was washed twice with 75% ice-cold ethanol, air dried for 20 min at room temperature and resuspended in 1x elution buffer. For both gDNA extractions we performed standard quality control, which involved Qubit quantification, Nanodrop measurement, and pulse-field gel electrophoresis making use of the Femto Pulse system (Agilent Technologies).

For *Idiurus macrotis*, *Desmana moschata*, *Muscardinus avellanarius*, *Cathorops nuchalis*, *Cathorops wayuu*, *Xerotyphlops vermicularis, Dipus sagitta*, *Ptilocercus lowii* and the two *Anguis fragilis* samples, gDNA was extracted according to the protocol of [49]. DNA concentration and DNA fragment length were assessed using the Qubit dsDNA BR Assay kit on the Qubit Fluorometer (Thermo Fisher Scientific) and the Genomic DNA Screen Tape on the Agilent 4150 TapeStation system (Agilent Technologies). For *Elysia timida* and *Scintilla philippinensis*, gDNA was extracted using a CTAB-based method [50] and a bead-based protocol [51], respectively, including a pre-wash with sorbitol. The MagAttract HMW DNA Kit from Qiagen was used to extract gDNA from *Podura aquatica.* For these gDNA extractions, DNA concentration and DNA fragment length were assessed using Qubit quantification (Thermo Fisher Scientific), the Agilent 2200 TapeStation system (Agilent Technologies) and the Femto Pulse system (Agilent Technologies).

All details on the DNA yield and DNA fragment sizes can be found in Supplementary Table 1.

### Low input PacBio HiFi library preparation

The low input protocol allows generating PacBio libraries for samples with limited DNA content without amplification [26]. We prepared low input PacBio HiFi libraries according to the instructions of the SMRTbell Express Prep Kit v2.0, except for the libraries of *Cathorops nuchalis* and *Cathorops wayuu* which were prepared with the SMRTbell prep kit v3.0.

### Ultra-low input PacBio HiFi library preparation

PacBio ultra-low input HiFi libraries were prepared with the SMRTbell Express Template Prep Kit 2.0 according to the ‘Procedure & Checklist - Preparing HiFi SMRTbell® Libraries from Ultra-Low DNA Input’ (PN 101-987-800 Version 02). To reduce potential PCR bias of polymerase A/B, we used in our modified protocol a third PCR reaction, making use of Polymerase C (KOD Xtreme™ Hot Start DNA Polymerase, Merck PN 71975), which is optimized for the amplification of long strands and GC-rich DNA templates.

The amplified DNA from two PCR reactions with polymerase A and B was pooled equimolarly. PCR fragments from polymerase C amplification were kept separately and processed independently from the pooled fragments produced with polymerase A and B. Purified and pooled amplified DNA libraries were size selected to remove smaller fragments (Supplementary Table 1).

For *Anguis fragilis* and *Idiurus macrotis*, we prepared two additional libraries with DNA extracts to which a DNA repair step was applied using the Sequential Reaction Protocol for PreCR Repair Mix (New England BioLabs) prior to the actual library preparation.

### PacBio sequencing

A total of 27 SMRT 8M cells were sequenced in CCS mode using the PacBio Sequel II / IIe instrument. For low input libraries, where possible, libraries were loaded at an on-plate concentration of 80 pM using adaptive loading and the Sequel II Binding kit 2.2 or 3.2 (Pacific Biosciences, Menlo Park, CA). Ultra-low input libraries were loaded with up to 80 pM on plate where possible using the SEQUEL II binding kit 2.2 or 3.2, and the sequencing kit 2.0. Pre-extension time was 2 hours, run time was 30 hours.

### HiC for scaffolding the *B. torquatus* assembly

Chromatin conformation capture was done using the Arima HiC+ Kit (Material Nr. A410110), following the user guide for animal tissues (ARIMA-HiC 2.0 kit Document Nr: A160162 v00) and processing 28 mg of tissue with the standard input approach. The subsequent Illumina library preparation followed the ARIMA user guide for Library preparation using the Kapa Hyper Prep kit (ARIMA Document Part Number A160139 v00). The barcoded HiC libraries were run on an S4 flow cell of a NovaSeq6000 with 200 cycles.

### Comparing polymerase A/B and C read assemblies

Aiming to evaluate the impact of libraries generated with polymerase A/B vs. C on the genome assembly quality, we combined different datasets with varying coverages, library complexities (number of libraries) and polymerase combinations (only A/B, only C, and A/B+C). For tests that did not involve all read data, we randomly subsampled reads. Subsequently, we assembled the read data into a contig assembly, as described below, and compared the summary metrics, including contig N50, number of contigs and gene completeness. All results are listed in Supplementary Tables 4 and 5.

### Contig assembly

HiFi reads were called using a pipeline consisting of PacBio’s tools ccs 6.4.0 (https://github.com/PacificBiosciences/ccs) and actc 0.3.1 (https://github.com/PacificBiosciences/actc) as well as samtools 1.15 [52] and DeepConsensus 0.2.0 or 1.2.0 [27]. All commands were executed as recommended in the respective guide for DeepConsensus (https://github.com/google/deepconsensus/blob/v0.2.0/docs/quick_start.md; e.g. ccs --all). To remove PCR adapters and PCR duplicates, which might originate from the PCR amplification during the ultra-low library preparation, PacBio’s tools lima 2.6.0 (https://github.com/PacificBiosciences/barcoding) with options “--num-threads 67 --split-bam-named --same” and pbmarkdup 1.0.2-0 with options “--num-threads 67 --log-level INFO --log-file pbmarkdup.log --cross-library --rmdup” (https://github.com/PacificBiosciences/pbmarkdup) were applied to samples prepared with the ultra-low library preparation protocol. For the Catfish samples *Cathorops nuchalis* and *C. wayuu* that were sequenced using the low-input library preparation protocol, PacBio sequencing adapters were removed with HiFiAdapterFilt [53]. The resulting reads were merged and then decontaminated with kraken2 v. 2.1.3 [54] using the kraken2 PlusPFP database downloaded in March 2023, with a confidence score of 0.51.

After HiFi calling, we used hifiasm v0.19.5 [28,55] to assemble HiFi reads obtained from the *Cathorops nuchalis, C. wayuu, Idiurus, Anguis, Elysia, Scintilla* and *Podura* samples. For the two catfish samples *Cathorops nuchalis* and *C. wayuu*, because of suboptimal performance with default parameters, we tested several hifiasm options before deciding which parameters produce the best assembly in terms of gene completeness and contiguity (Supplementary Table 2). To this end, we estimated the genome profile of these two species with FastK (https://github.com/thegenemyers/FASTK) and Genescope.FK (https://github.com/thegenemyers/GENESCOPE.FK) with k=30 to find the homozygous peak that was then passed to hifiasm (Supplementary Table 2). In all other cases, we applied default parameters with strict haplotig purging (-l3 parameter), and for the *Elysia* sample, we additionally used available Arima HiC data for assembly phasing.

Contiguity statistics were calculated with Quast 5.0.2 [56], gfastats v. 1.3.6 (https://github.com/vgl-hub/gfastats) and Merqury.FK (https://github.com/thegenemyers/MERQURY.FK). Gene completeness was evaluated with BUSCO 5.5.0 [30] as well as compleasm 0.2.5 [29]. We used the eutherian_odb10 dataset for *Bradypus torquatus*, and actinopterygii_odb10 for *C. nuchalis* and *C. wayuu*, the mammalia_odb10 dataset for human, the arthropoda_odb10 dataset for *Podura aquatica,* and the metazoa_odb10 dataset for *Elysia timida* and *Scintilla philippinensis*.

For *B. torquatus*, we initially obtained hifiasm (v0.19.5) assemblies that were of a size expected from four haplotypes of this genome, consisting of a large number of small contigs (Supplementary Table 6). Similar results were obtained with HiCanu (v2.2) [57], which is designed to break contigs at all joins in the assembly graph, meaning any divergences between the four theoretical haplotypes would result in a new contig (in our case over 200,000 assembled contigs totaling almost 12 Gb of sequence). This indicated that the tissue samples we obtained for this species originated from two different individuals. While the accuracy of the PacBio HiFi reads should in principle allow to distinguish all four haplotypes, *B. torquatus* is expected to have a very low heterozygosity and high in-breeding rate due to small population size, which results in assembly graphs where many regions collapse all haplotypes due to the lack of sequence variation.

To overcome this problem, we used the assembler Flye (v2.9.2) [58], which allows users to set the read error rate as an argument. Flye has been previously suggested by the developers as a method for collapsing sequences from highly diverged haplotypes into a single “pseudo-haplotype” sequence (https://github.com/fenderglass/Flye/issues/636). Here, we found that a read error-rate of 3% produced the most contiguous assembly, when combined with a reduced read-overlap of 5 kb (Supplementary Table 6). The latter deviates from the default value selected by Flye, which Flye would determine by the N90 of the input reads (in our case the N90 was 9 kb for the HiFi library, which had a modal read length of ∼10 kb). We then removed retained haplotigs using purge-dups [59].

### Contamination detection and read coverage analysis

Specimens stored in liquid preservation media are prone to various levels of DNA contamination from non-target organisms [33], caused by different handling and storage conditions that are often hard to retrace [60]. To detect levels of contamination from exogenous DNA in our assemblies, we used NCBI’s Foreign Contamination Screen (FCS 0.5.0) [43], which flags both putative adapter sequences (FCS-adaptor) and contigs assigned to non-target species (FCS-GX). Both FCS tools were executed from the provided singularity container using singularity 1.2.4. FCS-adaptor was executed through the provided bash script (run_fcsadaptor.sh) with the option for eukaryotes (--euk). FCS-GX was executed by the python wrapper (fcs.py screen genome) together with the corresponding NCBI taxonomy ID and the GX database (as of Dec 5th, 2023). Furthermore, to visualize contamination across the respective contig-level assemblies before FCS-filtering, we used blobtoolkit v4.1.4 [44], which assigns all contigs from a given assembly to a taxonomic group based on best blast hits (Supplementary Figure 2).

Additionally, to assess pre-assembly read quality, we mapped reads obtained from samples of *Bradypus torquatus*, *Anguis fragilis* and *Idiurus macrotis* to available reference genomes of closely related species *Choloepus didactylus* (GCA_015220235.1), *Elgaria multicarinata* (GCA_023053635.1) and *Pedetes capensis* (GCA_007922755.1). Similarly, to identify regions of PCR coverage dropouts, we aligned reads from polymerase A/B or C libraries to the best (defined as highest contig N50, Supplementary Table 5) assemblies obtained for *Podura*, *Scintilla* and *Elysia*, and visually inspected mapped reads (Supplementary Figures 5, 6).

To further quantify PCR bias, we calculated the normalized coverage (coverage of each nucleotide divided by the average coverage) of each polymerase A/B and C *Bradypus torquatus* library, using either the *Choloepus didactylus* genome or the best assembly of *Bradypus torquatus*. We also calculated normalized coverage of a non-amplified human library (downloaded from https://downloads.pacbcloud.com/public/revio/2022Q4/HG002-rep1/; last accessed 19 Sep 2024) as well as polymerase C (produced in this study) and A/B amplified libraries (NCBI, BioProject PRJNA657245, accessions SRR12454519 and SRR12454520) sequenced from the human cell line HG002, using the human HG002 assembly [6] (v.1.1, maternal haplotype). We then computed normalized coverage across nucleotides assigned to exonic and repeat sequences. For *C. didactylus* and *B. torquatus*, exons were annotated by TOGA v1.0.0 and repeats were annotated with RepeatModeler [61] and RepeatMasker v4.1.4 (https://www.repeatmasker.org/). For human, exons were annotated by RefSeq (v110 from CHM13, JHU v5.2), https://ccb.jhu.edu/T2T.shtml) and annotated repeats [62] were downloaded from the UCSC table browser [63] (last accessed 19 Sep 2024). Read mapping was performed using minimap2 v2.26 [64] with HiFi read mapping parameters (--ax map-hifi), and absolute coverage per base and across annotations was computed with samtools v1.17 [52], using the ‘samtools depth’ and ‘samtools bedcov’ commands, respectively. For the HG002 gene annotation, we filtered the annotation to only include coding exons to enable a fair comparison with the TOGA annotations that do not include non-coding transcripts or UTRs.

### Scaffolding the final *B. torquatus* genome

To scaffold these *B. torquatus* contigs, we mapped HiC reads to the contig assembly using bwa-mem (v.0.7.17) [65], before the resulting HiC alignment file was filtered, sorted and deduplicated with pairtools parse, pairtools sort and pairtools dedup (v0.3.0), respectively. The processed HiC alignments were then used as input for scaffolder yahs (v1.2a.1.patch) [39]. A full list of commands is given in Supplementary Note 1. After initial automated scaffolding with yahs, we ran multiple rounds of manual curation based on the HiC interaction maps. This involved re-ordering and re-orienting the scaffolded sequences based on sequences close to each other in the genome, which are expected to have a higher number of HiC interactions than those further apart. Using this method, we were able to obtain chromosome-level scaffolds of the 24 autosomes and the X chromosome. This assembly was then again screened for adapter and foreign sequence contaminants using NCBI’s FCS-adaptor and FCS-GX tools [43]. We subsequently removed contaminant sequences by applying the python wrapper (fcs.py clean genome) together with the action report from “screen genome” and setting the minimum sequence length to 1 bp (--min-seq-len 1).

### Read chimer analysis

To investigate whether our modified amplification based protocol creates more chimeric reads, we mapped reads (all obtained from the human HG002 sample) against the HG002 reference genome [6] (v.1.1, maternal haplotype, https://github.com/marbl/hg002?tab=readme-ov-file), using minimap2 v2.26 [64] with HiFi read mapping parameters (--ax map-hifi). We used reads amplified with polymerase C and polymerase A/B (NCBI BioProject PRJNA657245, accessions SRR12454519 and SRR12454520), as well as the non-amplified reads (https://downloads.pacbcloud.com/public/revio/2022Q4/HG002-rep1/; last accessed 19 Sep 2024), and reads amplified with MDA (NCBI BioProject PRJNA1005794, accession SRR25653511). To calculate the fraction of alignments classified as primary alignments, secondary alignments, supplementary alignments and unmapped, we counted the flags assigned by minimap using samtools v1.17 [52] with the command ‘samtools view’. Raw read lengths and alignment lengths of primary and supplementary alignments were extracted from raw fastq-files and sam-files created by minimap2, respectively.

## Acknowledgment

We thank Deniz Kaya from PacBio for advice and suggestions on adapting the ultra-low input protocol, and Sarah Kingan, Juniper Lake, Ian McLaughlin, Aaron Wenger and Jonas Korlach from PacBio for the polymerase C runs for the human sample. We also thank the Genome Technology Center (RGTC) at Radboudumc for the use of the Sequencing Core Facility (Nijmegen, The Netherlands), which provided the PacBio SMRT sequencing service on the Sequel IIe platform, the Long Read Team of the DRESDEN Concept Genome Center, part of the MPI-CBG and the technology platform of the CMCB at the TU Dresden, supported by DFG (INST 269/768-1), and the HPC Service of FUB-IT, Freie Universität Berlin, for computing time (doi:10.17169/refubium-26754). We acknowledge Irina Ruf (Senckenberg Frankfurt), Carles Galià Camps (University of Barcelona), Madlen Stange, Claudia Koch, Morris Flecks, Jan Decher and Christian Montermann (Leibniz Institute for the Analysis of Biodiversity Change) for providing samples and Sandra Kukowka for helping in subsampling specimens.

## Data and Code Availability

The raw sequencing data and assemblies for *Bradypus torquatus* are available at NCBI under BioProject PRJEB73341 and BioSample SAMEA115348596. An improved version of the *Elysia timida* assembly after incorporating additional polymerase C reads and Hi-C scaffolding [66] is available under Bioproject PRJNA1119176 and Biosample SAMN42332041. The *Scintilla philippinensis* assembly and raw sequencing data are available under Bioproject PRJNA1120792. Genome assemblies and raw sequencing data of both catfish genomes are available under Bioproject PRJNA1162287 (*Cathorops nuchalis*) and PRJNA1162286 (*Cathorops wayuu*). The *Podura aquatica* assembly and sequencing data are available under Bioproject PRJNA1163304. Raw reads and assemblies obtained with polymerase C for HG002 are available on https://downloads.pacbcloud.com/public/revio/2023Q3/KODXtreme/. The TOGA annotation for the *B. torquatus* is available at https://genome.senckenberg.de/download/TOGA/. Ultra-low input based assemblies generated in this study are also available at https://genome.senckenberg.de/download/GenomesCollectionsPolC. No new computer code was generated in this study.

