## Supplementary Figures for "Long-read sequencing and genome assembly of natural history collection samples and challenging specimens"

The Supplement contains:

- Supplementary Figures 1-6
- Supplementary Note 1

Supplementary Tables 1-6 are provided as sheets in a separate Excel file.

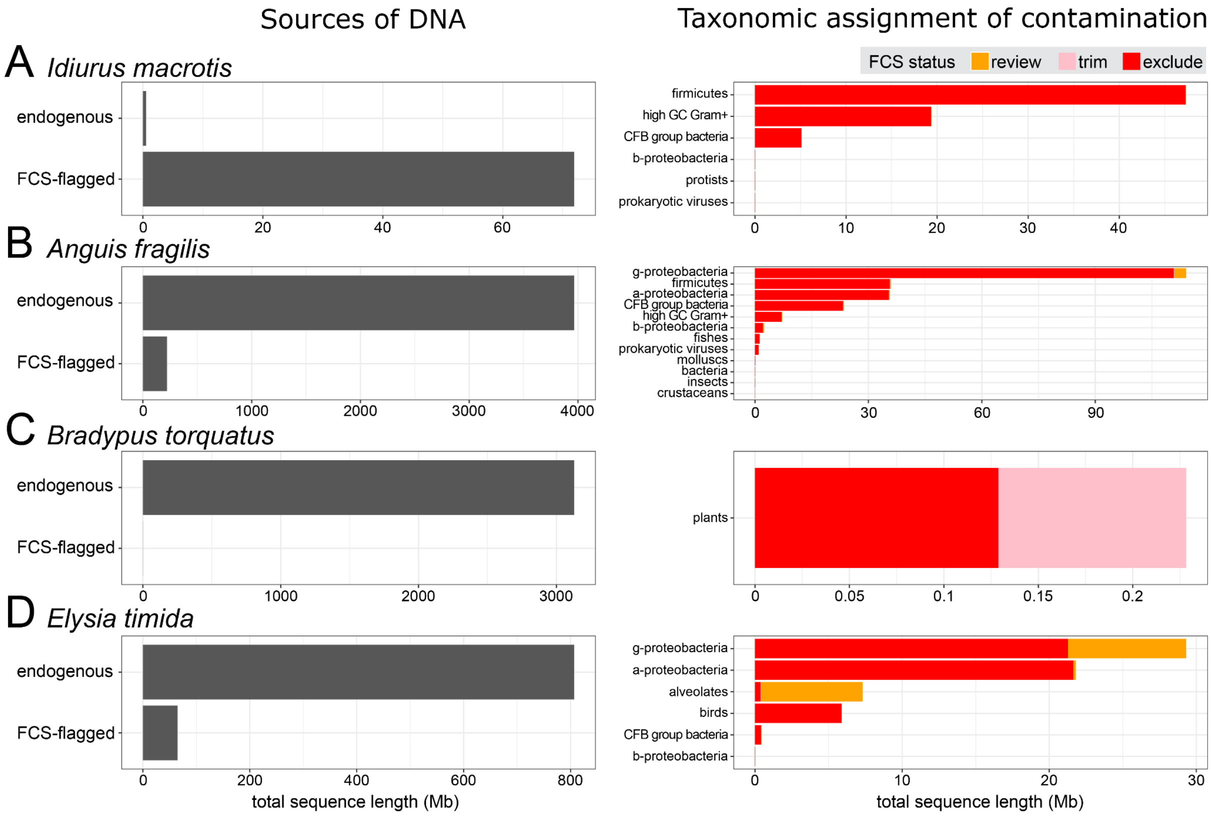


**Supplementary Figure 1: Levels of exogenous DNA contamination.**

For the samples sequenced with the PacBio ultra-low library protocol, we assembled the HiFi reads and screened the contigs for exogenous DNA contamination. The left plots show the total length of sequences not flagged (endogenous) or flagged (fcs-flagged) as contamination by FCS. The right plots show the total length of sequences assigned to a certain contaminant taxa. Sequences tagged by the FCS status “exclude” and “trim” show clear non-endogenous taxonomic signals, while sequences tagged with “review” are borderline cases.

(A) For contig assembly of reads obtained from the *Idiurus macrotis* sample, most of the sequenced DNA is contamination from bacteria. Aligning the sequenced reads to the genome of a close relative (*Pedetes capensis*) revealed an extremely low mapping rate of <0.1%, confirming that essentially no endogenous DNA is in the sample.

(B) For contig assembly of reads obtained from the *Anguis fragilis* sample from 2021, the majority of the assembled sequence is putatively endogenous DNA; however bacterial contamination is also present. Aligning the sequenced reads to the genome of a close relative (*Elgaria multicarinata*) showed a moderate mapping rate of 36,27%, confirming that a part of the sequenced DNA is endogenous.

(C) For the final chromosome-level assembly of the maned sloth (*Bradypus torquatus*), no contamination was found. FCS identifies very low levels of plant contamination. Consistent with these results for the final assembly, screening the HiFi reads of the first SMRT cell with kraken2 [1] indicated very low levels (2.6%) of non-mammalian DNA in the sample.

(D) The *Elysia timida* hifiasm-phased assembly of haplotype 1, generated from HiFi data obtained with both polymerases A/B and C, exhibits low levels of contamination mostly coming from bacteria.

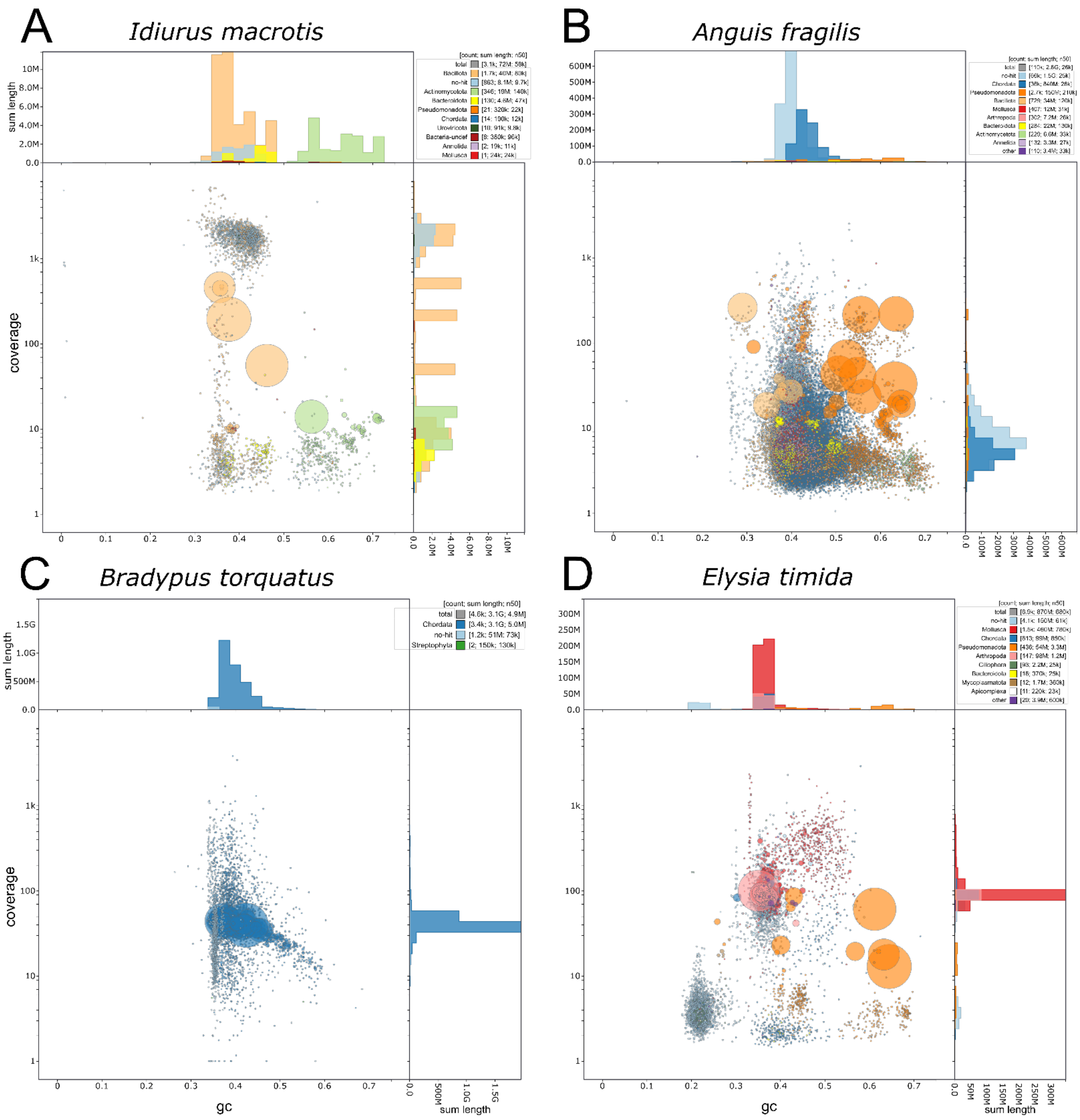


**Supplementary Figure 2: Breakdown of taxa contributing to exogenous DNA contamination.**

Blobplots created with blobtoolkit [2] for assemblies created with the ultra-low input protocol. The central plot depicts assembly contigs as blobs, with the blob sizes corresponding to contig sizes (scaled to the largest contig in the assembly) and colors representing the assigned taxon as inferred from the best blast hits. The Y-axis shows the coverage of sequenced reads mapped back to the respective contig and the X-axis shows the average GC-content of each contig. Histograms on top and right hand side summarize average GC-content and coverage of assembly contigs, partitioned by the assigned taxa.

(A) The contig assembly of *Idiurus macrotis* reads shows that most of the assembled contigs are assigned to different bacteria, and coverage as well as GC content of assembled contigs varies widely, indicating that the assembly mostly consists of contaminant sequence.

(B) For the contig assembly of the *Anguis fragilis* sample from 2021, the majority of the assembly is assigned to either "Chordata" (dark blue) or "no-hit" (light-blue), and "no-hit" contigs partly resemble "Chordata" contigs in terms of coverage and GC-content. On the other hand, the longest contigs in this assembly are assigned to “Pseudomonadota” (Proteobacteria, orange), meaning that there is evident bacterial contamination present in the assembly. This sample was collected as a roadkill, and partial decomposition by bacteria provides an explanation for the observed contamination.

(C) In the final contig level assembly of *Bradypus torquatus*, only two small contigs are assigned to “Streptopytha” (green), highlighting the absence of systematic contamination.

(D) The majority of contigs of the *Elysia timida* hifiasm-phased assembly of haplotype 1 is assigned to “Mollusca” (red), but the blobplot clearly shows several other clusters consisting of large contigs assigned to “Pseudomonadota” (orange) and other bacterial taxa or “no-hit” (light-blue). As the individual sequenced here was immediately killed and homogenized for sequencing without prior fixation, sample decomposition is unlikely. Rather, *Elysia timida* is too small to dissect before sequencing, therefore the observed contamination is likely caused by the gut microbiome and other commensal microorganisms. It should be noted that mollusca are underrepresented in the blast database, which likely increases false-positive hits and “no-hit” assignments [3].

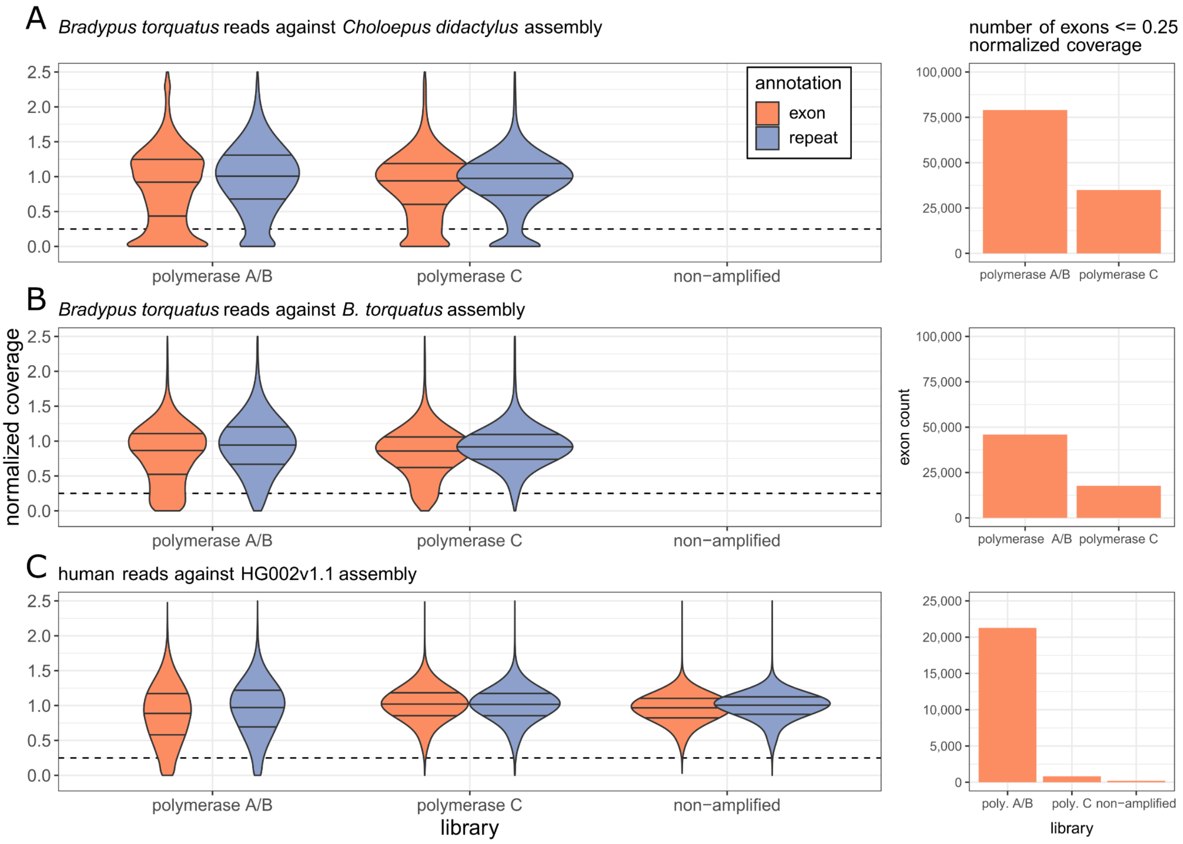


**Supplementary Figure 3: Amplification with polymerase C creates more even read coverage across exons and repeats.**

(Left) Violin plots depict the distribution density of normalized read coverage (Y-axis), calculated as the read coverage at a given genomic position divided by the average coverage of the respective library. Horizontal lines show 25% quartile boundaries, median and 75% quartile boundaries, respectively. Dotted lines represent a normalized coverage of 0.25. Only genomic regions annotated as exons or repeats were considered. Reads from polymerase C cover exonic and repeat regions more evenly, as the mode of the distribution is more pronounced around ~1.0 normalized coverage.

(Right) Number of unique exons having ≤0.25 normalized coverage. The overall number of unique exons is 256,011 (*Choloepus didactylus*), 246,006 (*Bradypus torquatus*), and 239,251 (human HG002). There is an up to ten-fold excess of exons having ≤0.25 coverage for polymerase A/B compared to polymerase C, highlighting potential drop-outs that can lead to a fragmented assembly and missing annotations.

(A,B) *Bradypus torquatus* sequencing reads obtained with polymerase A/B or C, aligned against the *Choloepus didactylus* genome (A) or the *Bradypus torquatus* genome (B).

(C) Human sequencing data aligned against the HG002v1.1 maternal haplotype assembly, comparing reads obtained with polymerase A/B or C and reads obtained without amplification.

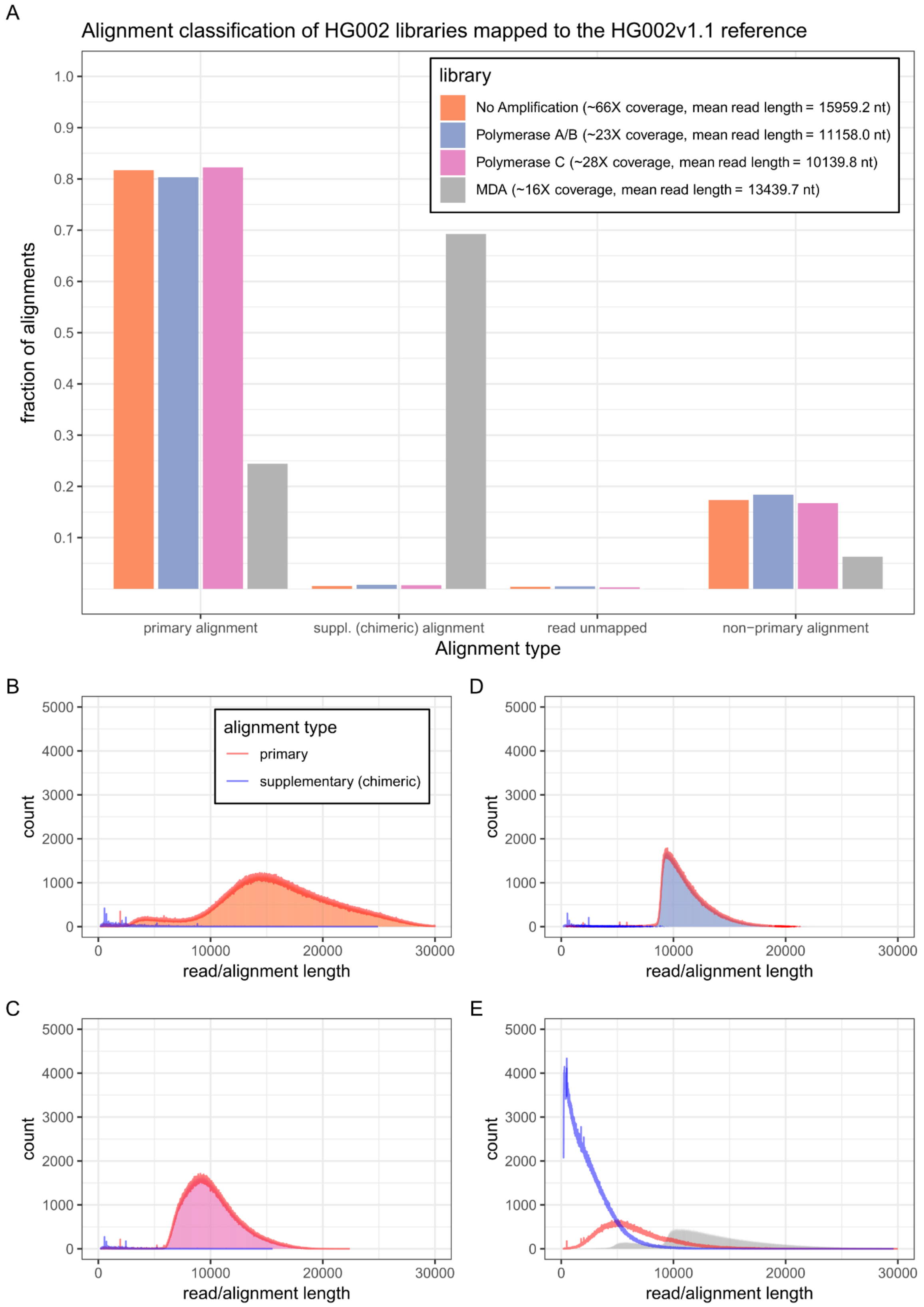


**Supplementary Figure 4: Low fraction of chimeric reads in amplified PacBio HiFi-read libraries.**

(A) The Y-axis shows different types of alignments as classified by minimap2 [4] when mapping human reads to the HG002v1.1 assembly. The X-axis depicts the fraction of alignments from a library. Supplementary alignments of a read indicate that the read is a chimer consisting of two different genomic regions. Alignments of non-amplified ultra-low input reads serve as a baseline and show a similarly low fraction of supplementary alignments (0.72%) as reads obtained with polymerase A/B (0.81%) and polymerase C (0.54%). In stark contrast, the majority of MDA reads have supplementary alignments and are likely chimeric (69.3%).

(B - E) Histograms of raw read and alignment lengths of different PacBio libraries, following the color scheme of A. The filled histogram areas indicate raw read length. The red and blue lines show primary and supplementary alignment lengths excluding soft- and hard-clipped bases. Whereas primary alignment lengths closely follow raw read length distributions for ultra-low and non-amplified reads (B,C,D), indicating full length mapping, for MDA reads (E), the mode of the primary alignment distribution is located at around ~5,000 nt compared to ~11,000 nt length for raw reads, clearly showing that primary alignments tend to be severely truncated. In accordance with (A), the amount and length of supplementary alignments is very low in non-amplified and ultra-low reads, but exceptionally high in MDA reads.

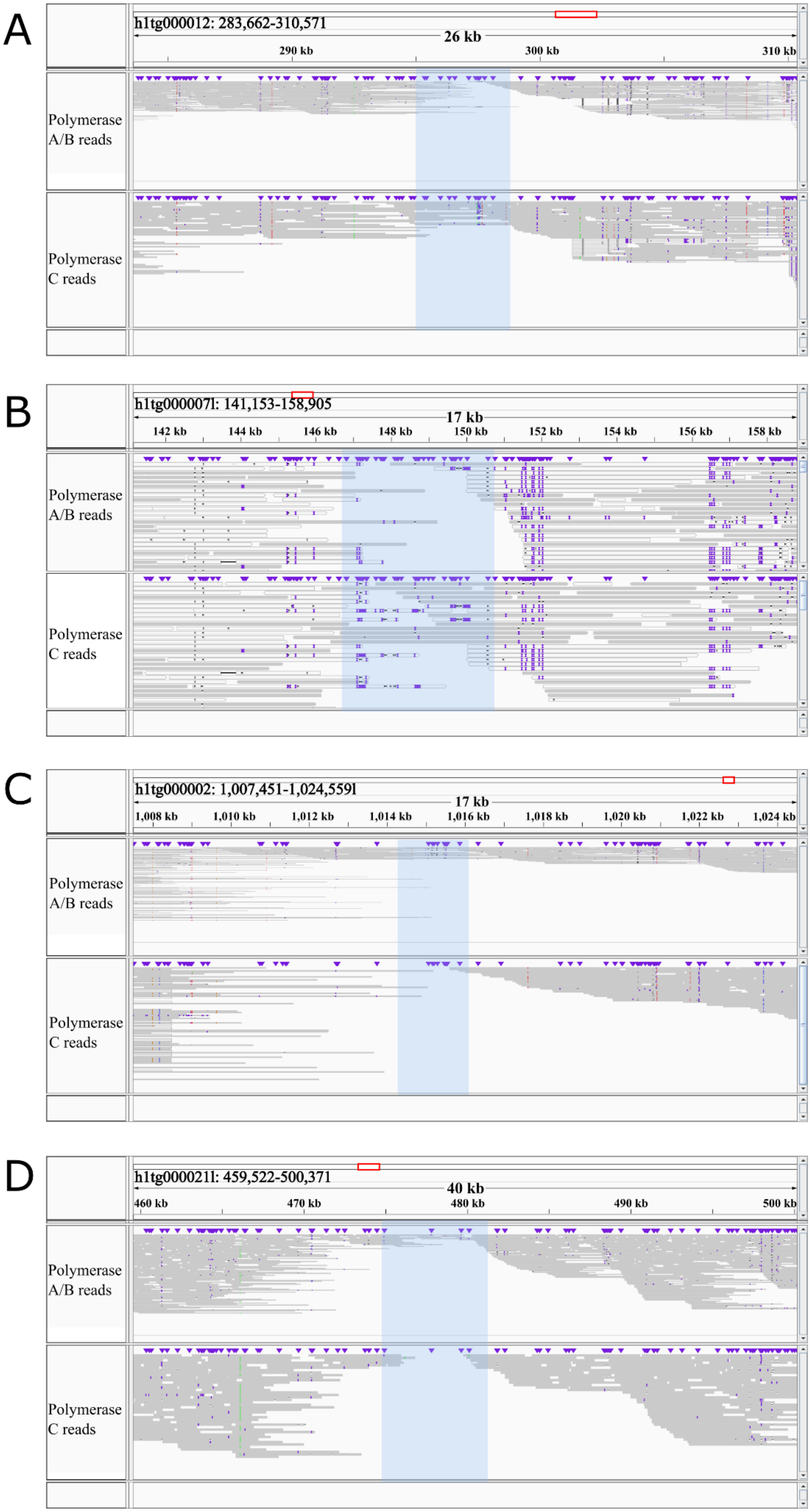
A
h1tg000012: 283,662-310,571
26 kb
290 kb
300 kb
310 kb
Polymerase A/B reads
Polymerase C reads
B
h1tg000071l: 141,153-158,905
17 kb
142 kb
144 kb
146 kb
148 kb
150 kb
152 kb
154 kb
156 kb
158 kb
Polymerase A/B reads
Polymerase C reads
C
h1tg000002: 1,007,451-1,024,559l
17 kb
1,008 kb
1,010 kb
1,012 kb
1,014 kb
1,016 kb
1,018 kb
1,020 kb
1,022 kb
1,024 kb
Polymerase A/B reads
Polymerase C reads
D
h1tg000211l: 459,522-500,371
40 kb
460 kb
470 kb
480 kb
490 kb
500 kb
Polymerase A/B reads
Polymerase C reads

**Supplementary Figure 5: Both polymerase A/B and C exhibit PCR bias for *Elysia timida*.**

IGV screenshots show alignments of HiFi reads produced with the ultra-low input protocol using either polymerase A/B or polymerase C. Reads were aligned to an assembly of *Elysia timida* generated with all reads from all three polymerases.

(A, B) Two examples of genomic regions where HiFi read coverage is very low or drops to zero for polymerase A/B while polymerase C reads cover the region. This exemplifies regions difficult to sequence with polymerase A/B.

(C, D) Two examples of loci where read coverage drops to zero for polymerase C but not polymerase A/B. While in (D) reads from polymerase A/B still cover the locus, it should be noted that there are fewer reads compared to the flanking regions. This indicates that it is also difficult for polymerase A/B to amplify these genomic regions.

#### *Scintilla philippinensis*

A

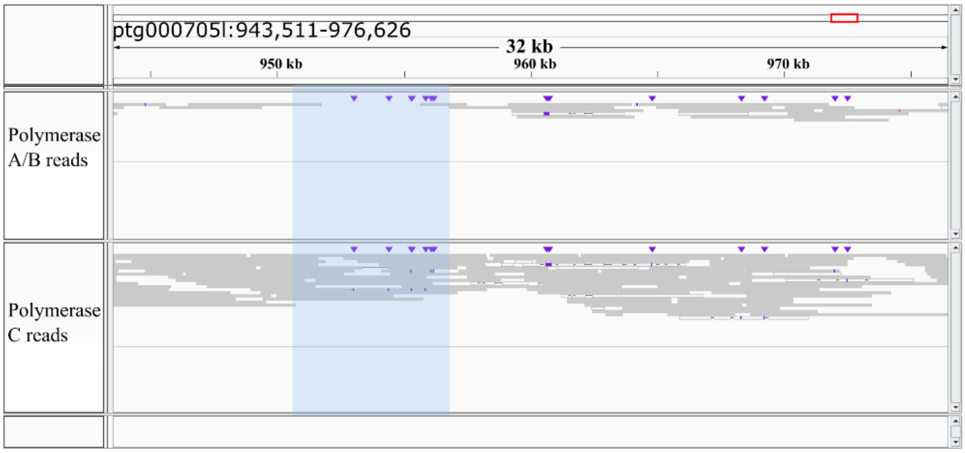


B

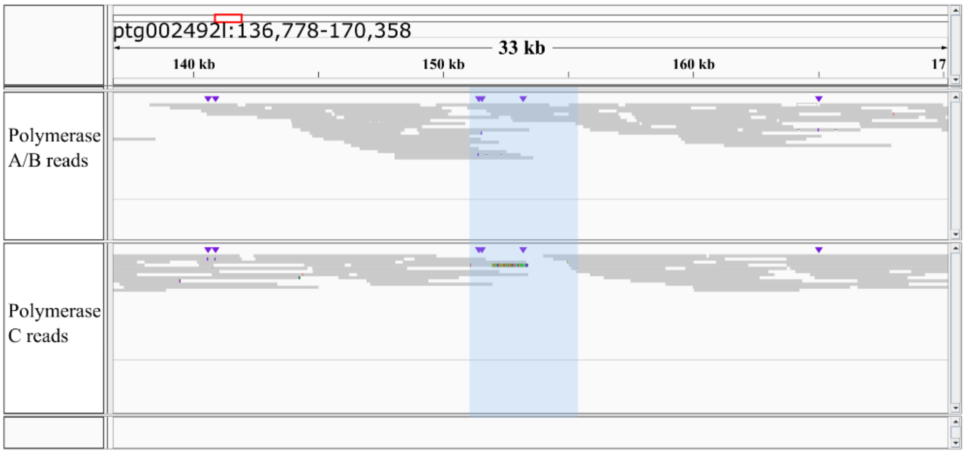


#### *Podura aquatica*

C

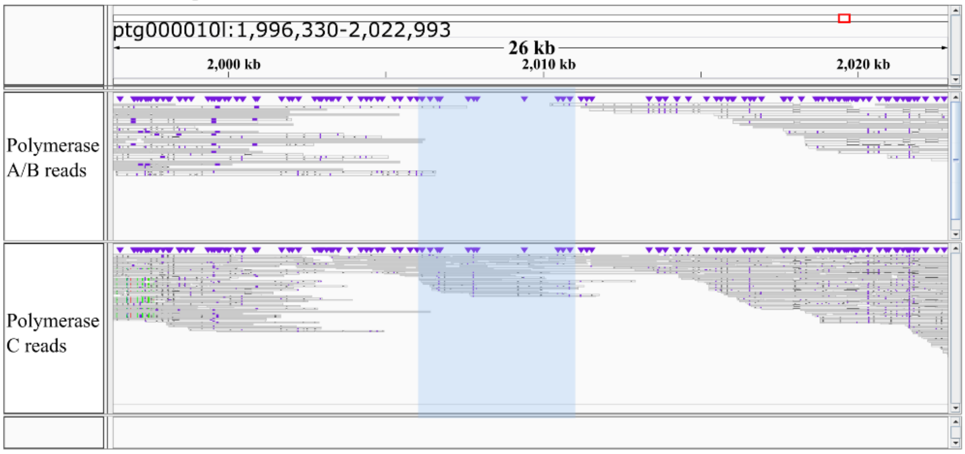


D

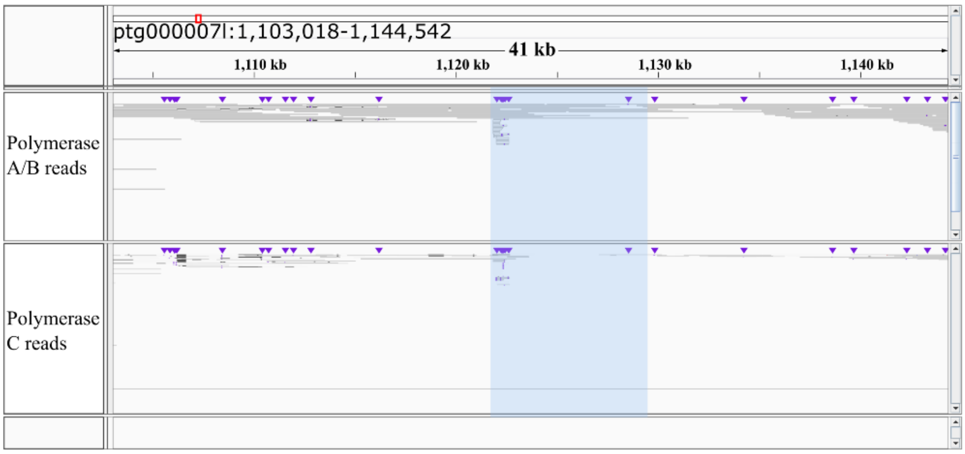

**Supplementary Figure 6: Polymerase A/B and C biases for *Podura aquatica* and *Scintilla philippinensis.***

IGV screenshots show alignments of HiFi reads produced with the ultra-low input protocol using either polymerase A/B or polymerase C. Reads were aligned to assemblies of *Scintilla philippinensis* (A, B) and *Podura aquatica* (C, D) generated with all reads from polymerases A/B and C.

(A, C) Two examples of genomic regions where HiFi read coverage is very low or drops to zero for polymerase A/B, while polymerase C reads cover these regions. This exemplifies regions difficult to sequence with polymerase A/B.

(B, D) Two examples of loci where read coverage drops to zero for polymerase C but not polymerase A/B. Of note, in panel D, polymerase A/B coverage is also comparatively low, and in panel B, there is a polymerase A/B coverage dropout upstream of the polymerase C dropout.

Together, this illustrates that for certain genomic regions one or both polymerases have difficulty amplifying DNA.

### Supplementary Note 1: Commands used for scaffolding the *Bradypus torquatus* assembly

```
bwa index asm_mBraTor.flye.purged.fa

bwa mem -5SP -T0 -t48 asm_mBraTor.flye.purged.fa <(zcat L122781_Track-
168234_R1.fastq.gz L122781_Track-168666_R1.fastq.gz) <(zcat L122781_Track-
168234_R2.fastq.gz L122781_Track-168666_R2.fastq.gz) -o mBraTor.bwa.sam

pairtools parse --output-stats mBraTor.parse.stats --min-mapq 40 --walks-policy 5unique --
max-inter-align-gap 30 --nproc-in 48 --nproc-out 48 --chroms-path
asm_mBraTor.flye.purged.fa.genome mBraTor.bwa.sam > mBraTor.bwa.parsed.pairsam

pairtools sort --nproc 48 --tmpdir=/scratch/brown/asm_mBraTor/assembly/yahs/
mBraTor.bwa.parsed.pairsam > mBraTor.bwa.sorted.pairsam

pairtools dedup --nproc-in 48 --nproc-out 48 --mark-dups --output-stats mBraTor.dedup.stats
--output mBraTor.bwa.dedup.pairsam mBraTor.bwa.sorted.pairsam

pairtools split --nproc-in 48 --nproc-out 48 --output-pairs mBraTor.bwa.dedup.pairs --output-
sam mBraTor.bwa.dedup.bam mBraTor.bwa.dedup.pairsam

samtools sort -@48 -n -T /scratch/brown/asm_mBraTor/assembly/yahs/ -o
mBraTor.bwa.dedup.sortname.bam mBraTor.bwa.dedup.bam
yahs-1.2a.1.patch/yahs asm_mBraTor.flye.purged.fa mBraTor.bwa.dedup.sortname.bam -v
1 -o mBraTor.yahs
```
